# mRNA translation control by Dhx36 binding to 5’UTR G-quadruplex structures is essential for skeletal muscle stem cell regenerative functions

**DOI:** 10.1101/2020.08.30.274068

**Authors:** Xiaona Chen, Jie Yuan, Guang Xue, Silvia Campanario Sanz, Di Wang, Wen Wang, Xi Mou, Mubarak Ishaq Umar, Joan Isern, Yu Zhao, Liangqiang He, Yuying Li, Christopher J. Mann, Xiaohua Yu, Lei Wang, Eusebio Perdiguero, Wei Chen, Yuanchao Xue, Yoshikuni Nagamine, Chun-Kit Kwok, Hao Sun, Pura Muñoz-Cánoves, Huating Wang

## Abstract

Skeletal muscle has a remarkable ability to regenerate owing to its resident stem cells, also called satellite cells (SCs), that are normally quiescent. When stimulated by damage, SCs activate and expand to form new fibers. The mechanisms underlying SC proliferative progression remain poorly understood. Here we show that Dhx36, a helicase that unwinds RNA quadruplex (rG4) structures, is essential for muscle regeneration by regulating SC expansion. We find that Dhx36 (initially named RHAU) is barely expressed at quiescence and is highly induced during SC activation and proliferation. Inducible deletion of *Dhx36* in adult SCs causes defective proliferation and muscle regeneration after damage. System-wide mapping in proliferating SCs revealed Dhx36 binding predominantly to rG4 structures at various regions of mRNAs, while integrated polysome profiling showed that Dhx36 promotes mRNA translation via 5’UTR rG4 binding. Furthermore, we demonstrate that Dhx36 specifically regulates the translation of *Gnai2* mRNA by unwinding its 5’UTR rG4 structures and identify Gnai2 as a downstream effector of Dhx36 for SC expansion. Altogether our findings uncover Dhx36 as an indispensable post-transcriptional regulator of SC function and muscle regeneration through binding and unwinding rG4 structures at 5’UTR of target mRNAs.

## Introduction

Skeletal muscle tissue homeostasis and regeneration relies on muscle stem cells, also known as satellite cells (SCs), which reside in a niche beneath the basal lamina attached to the myofibers, and are uniquely labeled by the expression of the paired box (Pax) transcription factor Pax7 [1–4]. SCs are typically in quiescence, a state of prolonged and reversible cell cycle arrest. Upon activation caused by injury or disease, SCs quickly re-enter the cell cycle and proliferate as myoblasts, a process orchestrated by the rapid induction of the master transcription factor (TF) MyoD. Subsequently, most myoblasts express Myogenin and differentiate, while a subset undergo self-renewal to restore the quiescent SC pool [1, 2]. Deregulated SC activity contributes to the progression of many muscle associated diseases [5]. At the cellular level, every phase of SC activity is tightly regulated by both intrinsic and niche-derived extrinsic factors [2, 6]. Elucidation of factors and molecular regulatory mechanisms governing SC function thus is of extreme importance, being the first step toward successful use of these cells in therapeutic strategies for muscle diseases.

RNA binding proteins (RBPs) regulate all aspects of RNA metabolism, including splicing and processing of mRNA-precursors (pre-mRNAs) in the nucleus, the export and localization of mRNAs to distinct subcellular regions in the cytoplasm, as well as mRNA translation and degradation [7]. Post-transcriptional regulation of gene expression by RBPs thus allows cells to orchestrate rapid changes in RNA or protein levels without altering transcription. Recent evidence suggests the contribution of post-transcriptional regulation to SC activities. For example, *MyoD* transcripts accumulate in quiescent SCs (QSCs), allowing rapid MyoD protein production as cells activate. High expression of Staufen1 in QSCs prevents *MyoD* mRNA translation by interacting with the secondary structure formed at its 3’UTR [8]. Additional studies showed that RBP-mediated RNA degradation plays key roles in SCs. In fact, proteins binding to AU-rich elements (ARE) located in 3’UTR of many mRNAs, such as AUF1[9], TTP[10] and HuR [11], regulate SC quiescence maintenance, activation and differentiation through modulating the stability of their interacting mRNAs. Still, molecular insights of SC post-transcriptional regulation remain largely unknown.

In this study, we discover new mechanisms of post-transcriptional regulation mediated by Dhx36, a DEAH-Box RNA and DNA helicase also known as RHAU or G4R1 [12, 13]. Dhx36 has emerged as a key RBP/helicase capable of binding and unwinding G-quadruplex (G4) structures, which are formed by guanine-rich nucleic acids harboring a motif [G_X_-N_1–7_-G_X_-N_1–7_-G_X_-N_1–7_-G_X_], where x is 3–6 nt and N corresponds to any nucleotide. These G4 structures typically consist of four tracts of guanines arranged in parallel or anti-parallel strands that align in stacked tetra planes, which are stabilized by Hoogsteen hydrogen bonds and a monovalent cation, and can be found in both DNAs and RNAs to provide additional layers of transcriptional or post-transcriptional regulation. RNA G4 (rG4) structures, in particular, are thought to participate in post-transcriptional regulation of mRNAs including pre-mRNA processing, mRNA turnover, targeting and translation [14–16], although the precise mechanisms are poorly understood. To date only a handful of proteins, including Dhx36, have been shown to bind and unwind rG4 *in vitro*. Dhx36 binds both DNA and RNA G4 structures with high affinity and specificity via a conserved N-terminal region known as the RHAU-specific motif (RSM) [17, 18]. Dhx36 is also capable of promoting mRNA translation by unwinding rG4s formed at 5’UTR (5’ untranslated region). For example, during cardiac development, Dhx36 binds to and unwinds the rG4 structure formed at the 5’UTR region of *Nkx2-5* mRNA, which is a key TF for heart development, to promote its translation [19]. Dhx36 can also exert post-transcriptional regulatory roles at other levels. In fact, Dhx36 was initially named RHAU (RNA helicase associated with AU-rich element) and shown to facilitate mRNA deadenylation and decay through direct association with the ARE element in *uPA* mRNA 3’ UTR, which lead to recruitment of PARN deadenylase and exosome [12]. Moreover, by binding and unwinding the rG4 formed in *pre-miR-26a*, Dhx36 promotes the maturation of *miR-26a* [20], and also resolves the rG4 formed in *p53* pre-mRNA, which is necessary for maintaining the 3’-end processing following UV-induced DNA damage [21]. Very recently, cross-linking immunoprecipitation sequencing (CLIP-seq) was used to profile DHX36 targets in Hela cells overexpressing DHX36, revealing that DHX36 preferentially interacts with G-rich and G4-forming sequences to increase the translational efficiency of target mRNAs [22]. As this study was based on Dhx36 overexpression it is not known whether the endogenous Dhx36 protein exhibits similar binding dynamics, and whether such binding profile exists in other cells. In particular, whether Dhx36 has a regulatory role in the mRNA metabolism of stem cells has never been addressed.

Dhx36 has a broad tissue expression and is indispensable for normal development. Accordingly, ablation of *Dhx36* in mouse is embryonic lethal and tissue-specific knock-out of *Dhx36* showed its requirement for hematopoiesis, spermatogonia differentiation and cardiac development [11–13]. However, its specific roles in somatic stem cells remain unexplored. In the present study, we investigated the function of Dhx36 in SC regenerative functions. By specifically inactivating *Dhx36* in mouse muscle SCs, we found that Dhx36 is required for normal muscle development and regeneration at adult age. Among the distinct myogenic stages, SC proliferation was particularly attenuated upon *Dhx36* deletion. Mechanistically, CLIP-seq showed that endogenous Dhx36 binds a large number of sites on thousands of mRNAs, being these binding sites G-rich and with high potential to form rG4 structures. Subsequent polysome profiling revealed that *Dhx36* loss leads to decreased translational efficiency of the target mRNAs with Dhx36 binding to their 5’UTR G4 sites, suggesting that the Dhx36-5’UTR G4 interaction functions to facilitate mRNA translation. Among the Dhx36-associated targets, we further confirmed the formation of rG4 at the *Gnai2* (Guanine nucleotide-binding protein G(i) subunit alpha-2) 5’UTR, with Dhx36 binding causing rG4 unwinding to facilitate *Gnai2* translation during SC activation and proliferation, thus identifying Gnai2 as a downstream effector of Dhx36 in regulating SC activity during muscle regeneration. Further in-depth analyses of the integrated CLIP-seq and polysome profiling datasets also revealed previously unknown aspects of Dhx36 binding and post-transcriptional regulation of mRNA processing. Altogether, our findings for the first time uncover the indispensable function of Dhx36 in muscle stem cells during adult muscle regeneration and provide a comprehensive mechanistic understanding of how this RNA helicase orchestrates post-transcriptional processes.

## Results

### Dhx36 is induced in activated SCs and required for normal muscle formation

To dissect how Dhx36 functions in SCs and muscle regeneration, we first examined *Dhx36* expression dynamics during SC myogenic progression by analyzing RNA-seq data. Quiescent SCs were isolated by fluorescence-activated cell sorting (FACS) from Pax7-nGFP mice [23], either fixed *in situ* by 0.5% paraformaldehyde (PFA) prior muscle digestion (SC_T0_) or from muscles digested without fixation (SC_T8_) (representing partially activated satellite cells by the 8 h isolation process) [24]. FACS-isolated SC_T8_ were further cultured for 24 h, 48 h or 72 h, representing fully activated or proliferating SCs (Fig. 1A). RNAs from the above cells were subject to RNA-seq. We found that the mRNA levels of *Dhx36* were very low in SC_T0_ and started to increase in SC_T8_, to peak at 48 h and remaining high at 72 h (Fig. 1B), thus suggesting that *Dhx36* expression is concomitant with the full activation and proliferation of SCs (Suppl. Fig. 1A). Western blot analysis of the Dhx36 protein confirmed that Dhx36 protein was induced in SCs cultured for 24 h and further increasing at 48 h (Fig. 1C). To further confirm the above expression dynamics of *Dhx36* in SCs *in vivo*, muscles of C57BL/6 mice were injected with cardiotoxin (CTX) to induce acute damage and regeneration. SCs are known to rapidly activate after muscle injury and reach a peak of proliferation at day 3, while they are mainly differentiating by day 7, coinciding with the initiation of myofiber repair [25, 26]. As expected, *Dhx36* mRNAs were highly enriched at day 3 (3 fold compared to uninjured) and started to decline by day 7 (Fig. 1D). To further examine its expression during the myogenic differentiation process *in vitro*, Dhx36 protein levels were also down-regulated when proliferating C2C12 mouse myoblasts were cultured in differentiating medium (DM), particularly at advanced differentiation stages (Suppl. Fig. 1B). Thus, Dhx36 expression is induced in activating and proliferating SCs, decreasing thereafter. Lastly, when examining its subcellular localization, we found that Dhx36 protein was exclusively present in the cytoplasm of activated SCs associated to visible puncta (Fig. 1E).

**Figure 1.**
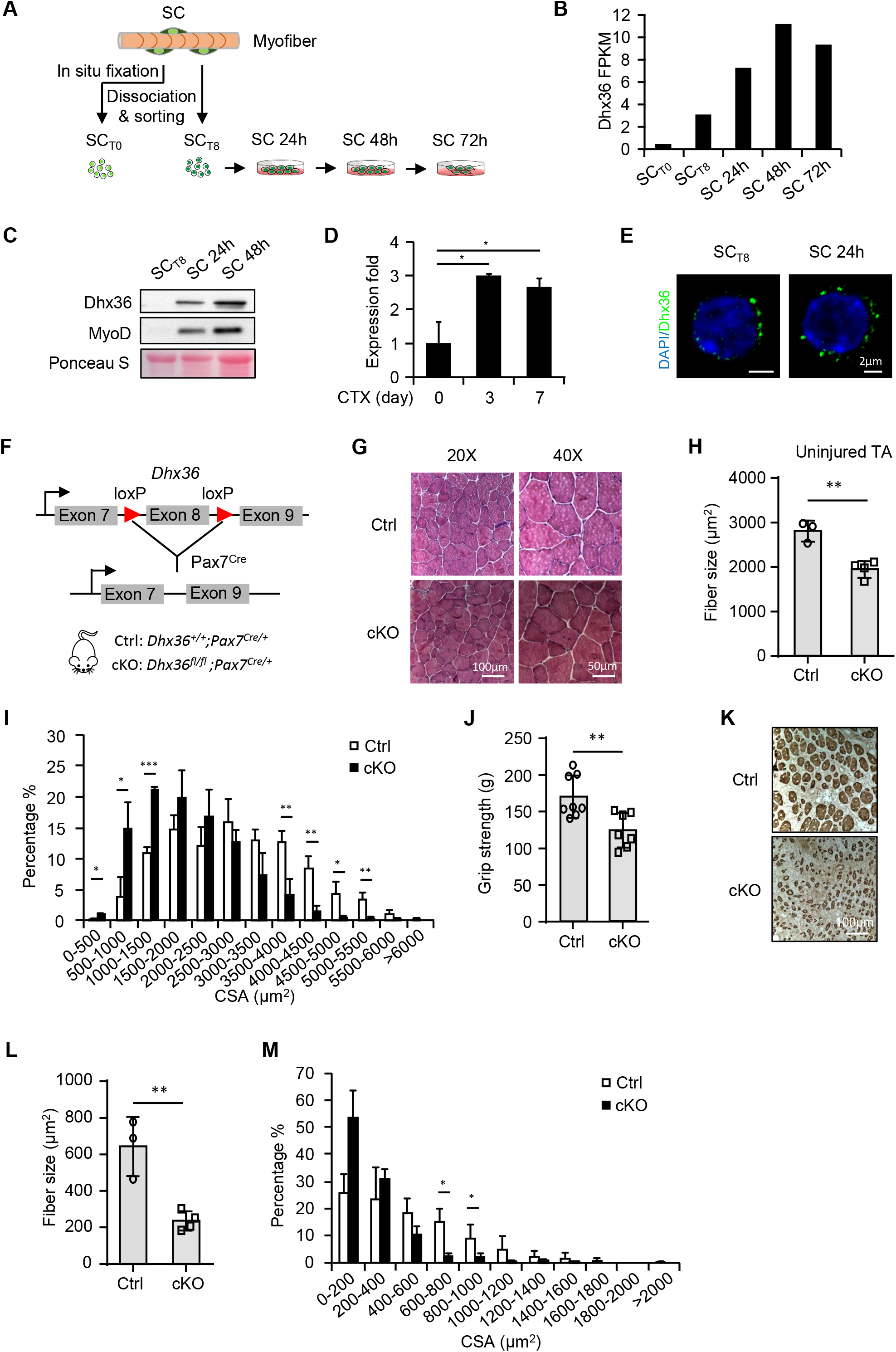
Dhx36 is induced in activated SCs and required for normal muscle development. (A) Schematic flowchart for isolation of quiescent satellite cells (SC_T0_) after in situ fixation with formaldehyde, or freshly isolated without prior fixation (SC_T8_) from Pax7-nGFP mice. SCs were cultured for 24 h, 48 h or 72 h to allow for activation and proliferation. (B) RNAs from the above cells were subject to RNA-seq and the expression dynamics (FPKM) of *Dhx36* mRNA is plotted. (C) Dhx36 protein levels in SC_T8_ or SC cultured for 24 h or 48 h were examined by Western blot. MyoD was used to indicate the activation of SCs. Total proteins on the membrane were stained by Ponceau S as equal loading control. (D) C57/BL-6 mice were injected with cardiotoxin (CTX) and SCs were isolated from uninjured muscle (day 0), or day 3 and 7 post-injection for qRT-PCR measurement of *Dhx36* mRNA expression. TATA-box binding protein (TBP) was used as normalization control. (E) Subcellular localization of Dhx36 protein in freshly isolated SCs or SCs cultured for 24 h was detected by immunofluorescent (IF) staining. High magnification confocal immunofluorescent representative images are shown. The green signal corresponds to Dhx36 immunoreactivity and nuclei are visualized by DAPI staining (blue). Scale bar: 2 μm. (F) Schematic illustration of breeding strategy to generate conditional knockout (cKO) mouse of *Dhx36*. *Dhx36^fl/fl^* mice were mated with *Pax7^Cre^* mice to inactivate *Dhx36* in cKO mice. Genotypes for Ctrl and *Dhx36* cKO mice are shown. (G) H&E staining of tibialis anterior (TA) muscle from adult Ctrl or *Dhx36* cKO mice. Scale bar=100μm (20x images) or 50μm (40x images). (H) Average fiber size in the above muscles was measured by ImageJ, Ctrl: n=3; cKO: n=4. (I) Cross section area (CSA) of each fiber from the above muscles was quantified and the distribution in each range is shown. (J) Grip strength of fore and hindlimb muscles were measured in Ctrl and cKO mice, Ctrl: n=8; cKO: n=7. (K) Immunohistochemisty (IHC) staining of eMyHC was performed on sections of TA muscles from Ctrl and cKO mice on day 7 post CTX induced muscle injury, scale bar=100μm. (L) The average fiber size of newly formed fibers was quantified from the above eMyHC+ fibers, Ctrl: n=3; cKO: n=4. (M) Distribution of CSA of newly formed fibers with central nuclei was quantified in TA muscles from Ctrl or cKO mice on day 7 post CTX induced muscle injury. Data represent the average of indicated No. of independent experiments□±□s.d. Student’s t-test (two-tailed unpaired) was used to calculate the statistical significance (D, H, I, J, L, M): *P < 0.05, **P < 0.01, ***P < 0.001.

Aiming to investigate the role of Dhx36 in skeletal muscle formation and regeneration, we first assessed whether constitutive deletion of *Dhx36* affects muscle formation. We conditionally deleted *Dhx36* in the Pax7 embryonic myogenic precursor cells by crossing *Pax7^Cre^* mice with *Dhx36^fl/fl^* mice in which the exon 8 of *Dhx36* was flanked by loxP sites [27–29] to generate conditional *Dhx36* knockout (cKO) mice (Fig. 1F). There was no evident defect of muscle development in cKO mice; adult body weight tended to be lower in cKO mice, although muscle/body weight ratio was comparable between Ctrl and cKO mice (Suppl. Fig. 1C-D). Morphometric analysis of tibialis anterior (TA) muscle revealed a decrease (30.7%) of average fiber size in cKO muscle compared with Ctrl muscle (Fig. 1G-H), and fiber size distribution revealed a higher number of smaller fibers in cKO mice (Fig. 1I). Moreover, the grip strength of fore and hind limb muscles of cKO mice was also decreased (22.6%) (Fig. 1J), suggesting an attenuated function of muscles lacking *Dhx36*. Finally, we examined the regenerative capacity of *Dhx36* cKO muscles, after injection of CTX to induce muscle damage and regeneration. Immunohistological staining of eMyHC (a marker protein for newly formed, regenerating myofibers) on cross sections of TA muscles harvested at day 7 post injury revealed a significant reduction of eMyHC+ fiber size (57.9%) in cKO muscles (Fig. 1K-M), indicating defective muscle regeneration in *Dhx36* mutant mice. Overall, these results imply that *Dhx36* loss in the early developmental SC precursors and progeny affects both normal growth and regeneration of skeletal muscle at adult age.

### Inducible loss of *Dhx36* in adult SCs impairs skeletal muscle regeneration

To further test *Dhx36* cell-autonomous functions in SCs during muscle regeneration, we crossed *Dhx36^fl/fl^* mice with *Pax7^CreER^/ROSA^EYFP^* mice (Fig. 2A) to specifically induce *Dhx36* deletion in adult SCs. Genetic inactivation of *Dhx36* in SCs was induced by intraperitoneal (IP) injection of tamoxifen (Tmx) in adult (8-12 weeks old) mice for five consecutive days followed by five days of chasing (Fig. 2B). Almost complete deletion of *Dhx36* was achieved in activated SCs of the inducible *Dhx36* knockout (iKO) compared to control (Ctrl) mice at both protein (Fig. 2C) and mRNA level (Suppl. Fig. 1E). No obvious morphological or histological muscle alterations were observed in iKO mice (data not shown).

**Figure 2.**
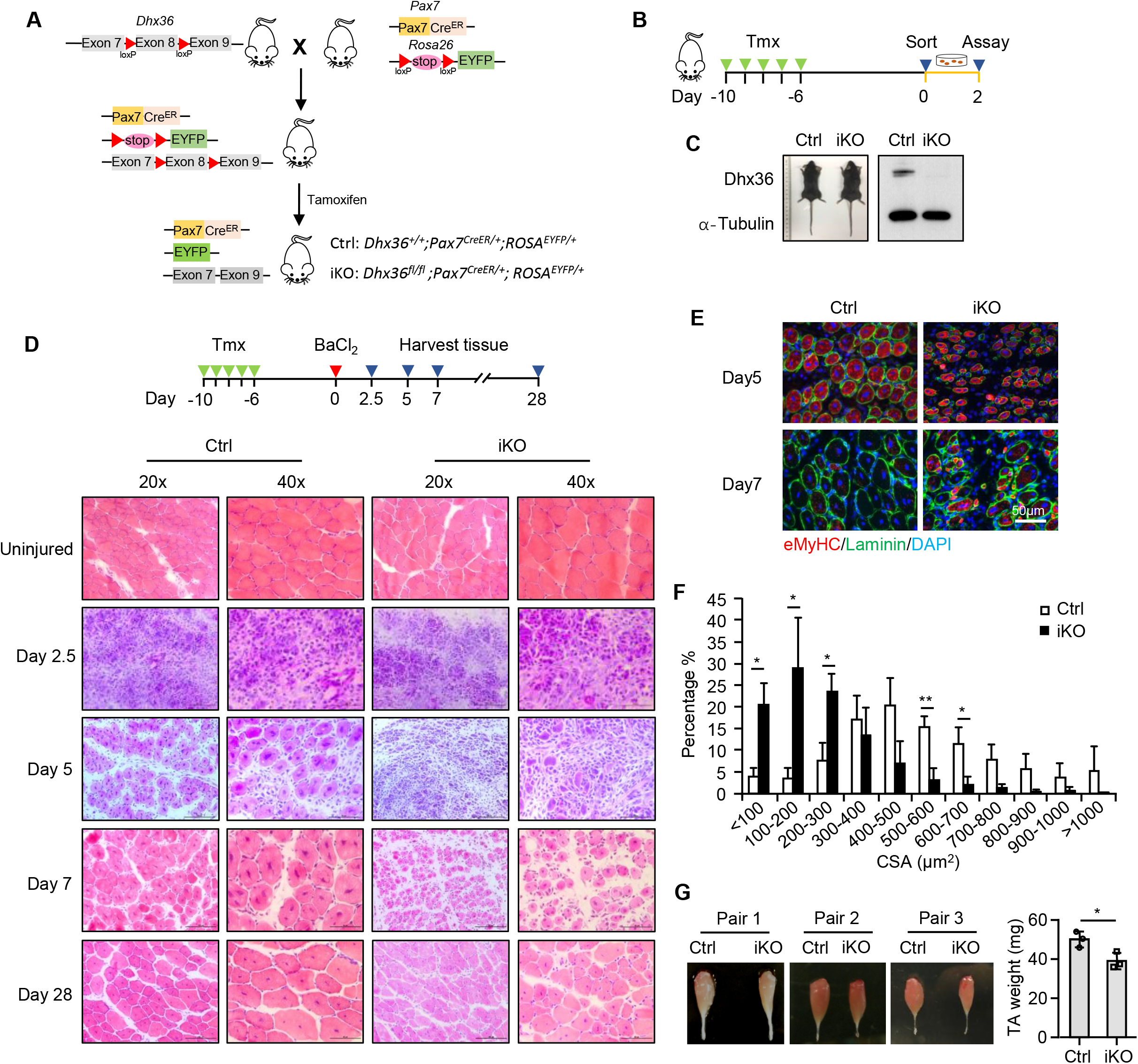
Inducible loss of *Dhx36* in adult SCs impairs skeletal muscle regeneration. (A) Schematic illustration of the strategy to inactivate *Dhx36* in inducible knock-out (iKO) mice. *Dhx36^fl/fl^* mice were mated with *Pax7^CreER^; ROSA^EYFP^* mice; the exon 8 of *Dhx36* was deleted in the iKO mice after Tamoxifen (Tmx) injection which also resulted in the removal of the stop signal for YFP at the Rosa26 site to allow the expression of YFP in iKO SCs. (B) Schematic illustration of 5 doses of Tmx injection to delete *Dhx36* in the iKO mouse. Freshly isolated SCs were collected at day 6 post-injection and cultured for two days. (C) Left: no obvious morphological difference was observed in representative Ctrl vs. iKO mice. Right: loss of Dhx36 protein was confirmed by Western blot with α-Tubulin as the loading control. (D) Upper panel: schematic illustration of the injury-induced muscle regeneration scheme. BaCl_2_ was injected into TA muscles of the above Ctrl or iKO mice 6 days post-Tmx injection to induce acute injury. The injected TA muscles were harvested at the designated times for the assessment of regeneration process. Lower panel: H&E staining of the TA muscles collected at 2.5, 5, 7 and 28 days post injury. Scale bar=100μm (20x) or50μm (40x). (E) IF staining of eMyHC (red) and laminin (green) was performed on the TA muscles collected at 5 and 7 days post BaCl_2_ injury. Nuclei are visualized by DAPI staining (blue). Scale bar=50μm. (F) CSAs of newly formed fibers were quantified from the above stained sections and the distribution is shown; n=3 mice per group. (G) Left: Representative images of TA muscles collected at 28 days post injury are shown. Right: the muscle weight from three pairs mice. Data represent the average of three independent experiments□±□s.d. Student’s t-test (two-tailed unpaired) was used to calculate the statistical significance (F, G): *P < 0.05, **P < 0.01.

We next examined whether loss of *Dhx36* in SCs impacts adult muscle regeneration, by intramuscular injection of BaCl_2_, which induces acute damage followed by regeneration to an extent similar to CTX injection, in Ctrl and iKO mice (Fig. 2D) [30]. Tissue degeneration with abundant immune cell infiltrates was observed both in iKO vs Ctrl muscles 2.5 days after injury. At day 5 after injury, regenerating fibers with centrally localized nuclei (CNL) were readily observed in Ctrl damaged muscles while they were rare in iKO damaged muscles, which presented still abundant immune cell infiltrates. By day 7 after injury, regeneration in iKO muscles was still significantly compromised, being the size of the newly formed myofibers much smaller than in Ctrl mice (Fig. 2D). eMyHC staining confirmed delayed muscle regeneration in iKO mice; 5 days after injury, both Ctrl and iKO injured muscles presented eMyHC+ fibers, being larger in the former mice as shown by myofiber cross-sectional area (CSA) measurement; at day 7 post injury, however, eMyHC+ fibers were no longer present in Ctrl muscles while they persisted in iKO muscles, consistent with the larger size of newly-formed myofibers in Ctrl muscles vs iKO muscles (Fig. 2E-F; Suppl. Fig. 1F). By day 28 post injury, regeneration of TA muscles advanced in both Ctrl and iKO mice; however, this process was still severely compromised in the mutant mice, as shown by the significantly smaller size of the new fibers in iKO compared to Ctrl muscles, consistent with a reduction (22.0%) of muscle weight (Fig. 2G). Altogether the above data demonstrated that deletion of *Dhx36* in SCs markedly blunts adult muscle regeneration, indicating its requirement for this process.

### *Dhx36* deletion principally impairs SC proliferative capacity

To pinpoint the defects of SCs upon *Dhx36* deletion, we investigated SC behavior *in vitro* and *in vivo*. First, we found that the number of Pax7+ SCs in Ctrl and iKO adult mice did not differ significantly 4 weeks after Tmx administration (Fig 3A), indicating *Dhx36* deletion does not impact the homeostatic SC quiescent state. Consistently, Pax7 staining also revealed no difference in SC number in single myofiber explants from iKO vs Ctrl mice (Suppl. Fig. 2A). To assess SC proliferative capacity, we measured EdU incorporation in SCs cultured for 2 days. As shown in Figure 3B, while 66.7% SCs from Ctrl mice were positive for EdU, only 39.7% were EdU+ in iKO mice, suggesting that the proliferative ability of SCs was compromised by *Dhx36* loss. These results were confirmed in single myofiber explants: the number of Pax7+ or YFP+ SCs on iKO myofibers was significantly lower compared with Ctrl (57.3% decrease for Pax7+ cells or 65.0% decrease for YFP+ cells) 2 days after isolation (Suppl. Fig. 2B and Fig. 3C). Moreover, in response to muscle damage, the number of Pax7+ SCs was also significantly decreased (52.9% decrease) in iKO vs Ctrl mice at day 3 post-injury (Fig. 3D). Finally, EdU incorporation in SCs *in vivo* after 2.5 days of muscle injury in Ctrl or iKO mice (all carrying Rosa EYFP reporter) showed that 31.8% of YFP+ cells in Ctrl mice were EdU+ while only 18.3% of YFP+ cells were EdU+ in iKO mice (Fig. 3G), confirming that loss of *Dhx36* in SCs compromises their proliferating capacity both *in vitro* and after muscle injury.

**Figure 3.**
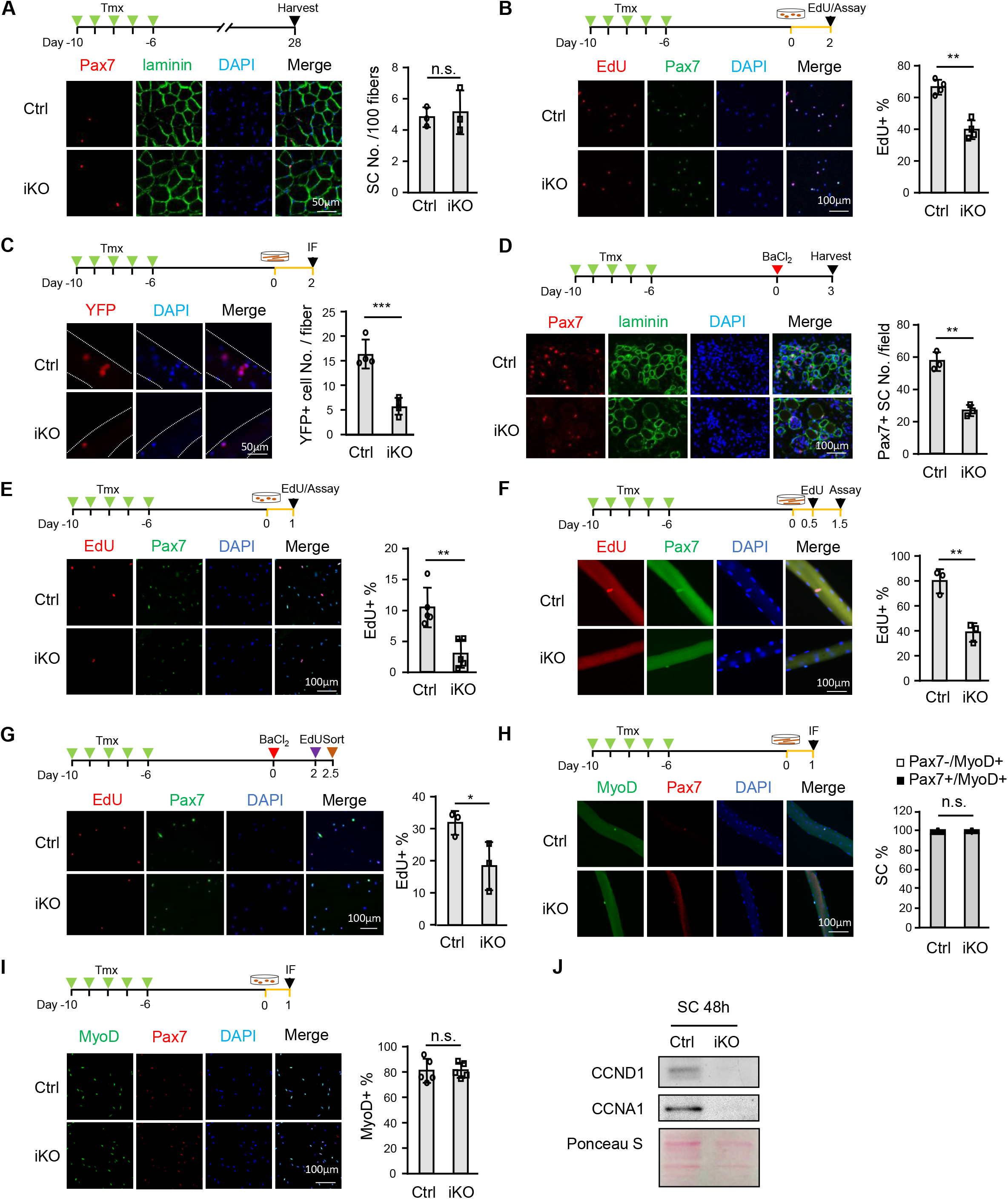
*Dhx36* deletion principally impairs SC proliferative capacity. (A) IF staining of Pax7 (red) and laminin (green) on cross-sections from Ctrl and iKO TA muscles collected at 4 weeks after 5 consecutive doses of Tmx injection. Cell nuclei were co-stained with DAPI (blue). Representative images are shown. The number of Pax7+ cells was quantified from >20 fibers/mouse. Scale bar=50μm; n=3 mice per group. (B) SCs from Ctrl or iKO mice were collected and cultured for 48h. EdU was then added to the culture medium 4 h before staining for EdU and Pax7. The percentage number of EdU+ cells was calculated from 10 randomly selected fields per mouse. Representative images are shown. Scale bar=100 μm; n=4 mice per group. (C) Single myofibers were isolated from Ctrl or iKO mice and cultured for 48 h. IF staining for YFP was performed and YFP+ cells was quantified from >15 fibers/mouse. Representative images are shown. Scale bar=50 μm; n=5 mice per group. (D) IF staining of Pax7 (red) and laminin (green) on sections of TA muscles at D3 post BaCl_2_ injection. Representative images are shown. Pax7+ cells numbers were calculated from 10 randomly selected fields per mouse. Scale bar=100 μm; n=3 mice per group. (E) SCs from Ctrl and iKO mice were cultured for 24 h with EdU and then fixed for EdU (red) and Pax7 (green) immunostaining. Representative images are shown. EdU incorporation percentage was calculated by counting from 10 randomly selected fields per mouse. Scale bar=100 μm; n=5 mice per group. (F) Isolated single myofibers were cultured with EdU for 36 hours and then fixed for immunostaining of EdU (red) and Pax7 (green). EdU incorporation percentage was calculated by counting from >15 fibers per mouse. Scale bar=100 μm; n=3 mice per group. (G) Lower hindlimb muscles were injured by BaCl_2_ followed by intraperitoneal injection of EdU at 48 h post-injury. Freshly isolated SCs from lower hindlimb muscles were collected 12 h later and stained for EdU (red). The nuclei were counterstained by DAPI (blue). The representative images are shown. EdU incorporation percentage was calculated by counting from 10 randomly selected fields per mouse. Scale bar=100 μm; n=3 mice per group. (H) IF staining of Pax7 (red) and MyoD (green) was performed on single myofibers cultured for 24 h. Percentages of Pax7+/MyoD+ and Pax7-/MyoD+ cells were quantified from >15 fibers per mouse. Scale bar=100 μm; n=3 mice per group. (I) IF staining of Pax7 (red) and MyoD (green) was performed on SCs cultured for 24 h. Percentages of MyoD+ cells were quantified from 10 randomly selected fields. Scale bar=100 μm; n=5 mice per group. (J) The protein levels of cyclin genes CCND1 and CCNA1 proteins were measured by Western blot in Ctrl and iKO SCs cultured for 48 h. Total proteins on the membrane were stained by Ponceau S as loading control. Data represent the average of indicated No. of independent experiments□±□s.d. Student’s t-test (two-tailed unpaired) was used to calculate the statistical significance (A-I): n.s., not significant, *P < 0.05, **P < 0.01, ***P < 0.001.

To further investigate whether the impaired proliferation of iKO SCs was caused by a defect in quiescence exit or the first cell cycle entry (*i.e.* the activation step), SCs isolated from Ctrl or iKO mice were cultured with EdU at early time points. 10.4% Ctrl SCs were EdU+ while only 3.0% iKO SCs incorporated EdU after 1 day (Fig. 3E); consistently, 38.7% of SCs on iKO fibers was EdU+ compared to 79.8% on Ctrl fibers after 36 h (Fig. 3F). Interestingly, co-staining of MyoD and Pax7 on single myofibers 24 h after isolation revealed that nearly all iKO SCs are MyoD+ showing no difference with Ctrl (Fig. 3H), suggesting the quiescence exit is not affected. Similarly, when freshly-sorted SCs were cultured for 1 day, a similar percentage of cells were MyoD positive in both Ctrl (81.0%) and iKO (81.4%) SCs (Fig. 3I) and there was no significant difference of MyoD expression at both mRNA and protein level between Ctrl and iKO SCs (Suppl. Fig. 2C-D). Moreover, a similar percentage of the above Ctrl (94.4%) and iKO (93.4%) SCs were Ki67+, suggesting no impairment in cell cycle re-entry but rather a defect in cell cycle progression (Suppl. Fig. 2E). To confirm the cell cycle progression defect upon *Dhx36* loss, we deleted *Dhx36* in C2C12 myoblasts by CRISPR-Cas9 mediated gene editing (Suppl. Fig. 2F). Indeed, homozygous deletion of *Dhx36* led to decreased EdU incorporation by 46% as compared with Ctrl clones (Suppl. Fig. 2G). Further cell cycle analysis by FACS revealed a severe G1 block in the KO cells after nocodazole treatment (Suppl. Fig. 2H); consistently, the cycling genes Ccnd1 and Ccna1 that regulate G1/S phase transition were also down-regulated in iKO vs Ctrl SCs (Fig. 3J). Altogether the above findings demonstrate that *Dhx36* is indispensable for timely activation and cell cycle progression of satellite cell-derived myoblasts during injury-induced muscle regeneration.

### Transcriptomic binding profiling uncovers diversified roles of Dhx36 in mRNA metabolism

The cytoplasmic localization in myoblasts (Fig. 1E) led us to speculate that *Dhx36* exerts its function as a post-transcriptional or translational regulator, consistent with its well-known rG4 and ARE binding and regulatory functions. Therefore, we conducted CLIP-seq in C2C12 myoblasts to map the RNA interactome of endogenous Dhx36 following the workflow shown in Suppl. Fig. 3A [33]. Using an antibody against Dhx36, we succeeded in retrieving endogenous Dhx36 at the expected size of ~115kd (Fig. 4A left) and the RNA-protein (RNP) complexes appeared 15-20 kD larger (Fig. 4A right). The bound RNA fragments were recovered from the RNPs and transformed into small RNA libraries for next generation sequencing. Through an in-house pipeline modified from [33], we identified a total of 5305 and 4982 binding sites from the two biological replicates which exhibited good reproducibility (Fig. 4B) and 3662 sites (corresponding to 1262 individual genes) were shared between the two replicates (Fig. 4C and Suppl. Table S1). Overall, the majority of the binding peaks (3032, 82.7%) were mapped to protein coding mRNAs and a small portion of bindings were found in non-coding RNA (ncRNA) (422), small nucleolar RNA (snoRNA) (142), or other RNA species (66) (Fig. 4D), indicating its predominant function in regulating mRNA processing. When examining the mRNA binding closely, we found a large number of binding sites resided in coding sequence (CDS) (2040) and introns (1345) (Fig. 4E); however, 5’UTR possessed the highest binding density (binding site number per 1000 nt) followed by CDS, 3’UTR and introns (Fig. 4F), suggesting its dominant functions through binding and regulating 5’UTRs. It is noteworthy that the majority of the target mRNAs possessed Dhx36 binding sites in multiple regions, for example, co-existing binding in 5’UTR, 3’UTR and CDS was found on 216 transcripts (Fig. 4G-H). Gene Ontology (GO) analysis of the 1262 genes associated with Dhx36 binding, showed enrichment in “regulation of gene expression, epigenetic”, “gene silencing”, and “DNA conformation change” (Fig. 4I), suggesting diversified gene regulatory functions of Dhx36.

**Figure 4.**
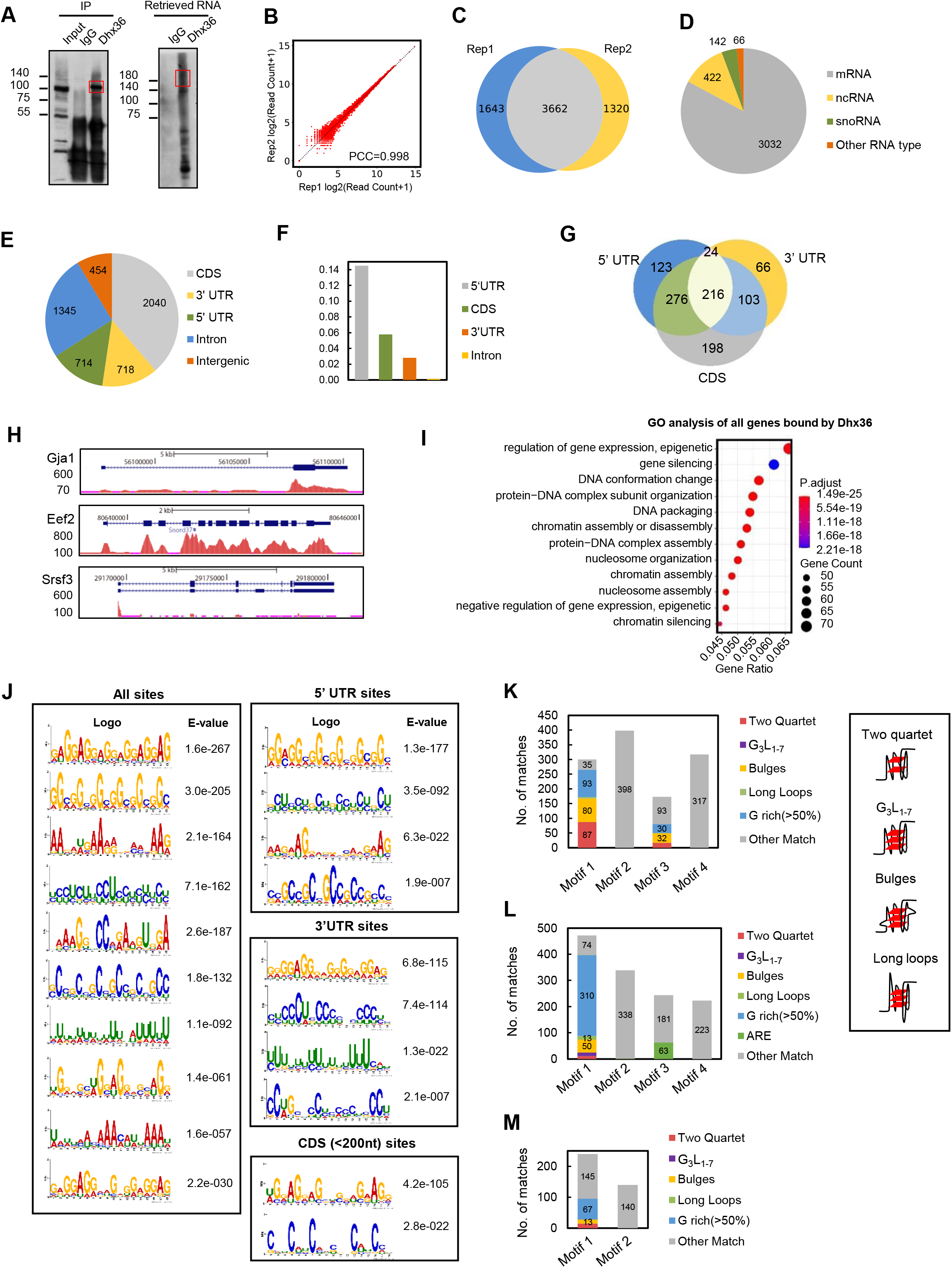
Transcriptomic binding profiling uncovers diversified roles of Dhx36 in mRNA metabolism. (A) CLIP-seq was performed in C2C12 myoblasts and Western blot shows an evident amount of Dhx36 protein was immunoprecipitated with an antibody against Dhx36 but not IgG. The above retrieved RNAs were labeled with biotin for the visualization and size selection. The indicated RNAs were recovered and used for sequencing. (B) Read coverages in the two CLIP-seq biological replicates displayed a high correlation. PCC, Pearson Correlation Coefficient. (C) A large number of identified binding sites (3662) were shared in the two biological replicates. (D) The categories of the Dhx36 associated RNAs are shown. (E) The distribution of the overlapped sites on different mRNA regions. (F) The density of Dhx36 binding sites in the designated regions of mRNAs. (G) Venn diagram showing the number of genes bound by Dhx36 in different mRNA regions. (H) Genome snapshots to illustrate Dhx36 binding in 3’UTR, CDS and 5’UTR on 3 selected transcripts. (I) The GO analysis results for all genes with Dhx36 binding ordered by gene ratio (proportion of genes annotated for each GO term). Dots are colored by adjusted P-value (degree of enrichment) and their size corresponds to the gene counts annotated to each GO term. (J) MEME identifies the top enriched motifs in all binding sites or sites residing in 5’UTRs, 3’UTRs or CDS. The E-values measuring the statistical significance of each motif identified by MEME are shown. (K) Potential rG4 formation was predicted in the top 4 enriched Dhx36 binding motifs in 5’UTR; the number of each subtype of rG4 (illustrated on the right) is shown. (L) Potential rG4 formation or ARE was predicted in the top 4 enriched Dhx36 binding motifs in 3’UTR; the number of each subtype of rG4 or ARE is shown. (M) Potential rG4 formation was predicted in the top 2 enriched Dhx36 binding motifs within CDS; the number of each subtype of rG4 is shown.

Further scanning of the binding motifs, using all the binding peaks by MEME [31], showed that G-rich sequences were highly enriched; G-rich sequences were also the top ranked when the scanning was performed in 5’UTR, 3’UTR or CDS binding sites alone (Fig. 4J and Suppl. Table S1). We next predicted whether these G-rich sequences were capable of forming rG4 structure using in-house python scripts (see Materials and Methods). Among the top four Dhx36 binding motifs enriched regions within 5’UTR, 57% of the top one motif were predicted to form G4 (Fig. 4K), highlighting that the primary function of Dhx36 in 5’UTRs is indeed related to rG4 binding. Interestingly, unlike previous reports showing Dhx36 binding with the canonical G4 sequence, G_3_L_1-7_ ((G_3_N_1-7_)_3_G_3_, N=A, U, C or G) G4 in 5’UTRs [32], bulges and two quartet appeared to dominate the predicted rG4s. In both 3’UTR and CDS regions, on the other hand, the formation of two quartet, G_3_L_1-7_, long loops and bulges rG4 was low (18% and 11.6%, respectively) (Fig. 4L-M). In addition to the G-rich sequences, one type of AREs (often defined as adjacent repeated AUUUA repeats) [33], was also found to be highly enriched (ranked No. 3) in the Dhx36 binding regions in 3’UTR (Fig. 4L) but not in 5’UTRs and CDS (Fig. 4J). Previously, Dhx36 was shown to bind with ARE motifs and regulate urokinase plasminogen activator (uPA) mRNA degradation [12]; our finding thus indicated that the binding to ARE may be a transcriptome-wide event that endows Dhx36 with key regulatory functions. Indeed, a total of 292 AREs were bound by Dhx36 corresponding to 154 genes and most of the sites resided in 3’UTR (Suppl. Fig. 3B and Suppl. Table S1); GO analysis revealed that the genes with Dhx36 binding in 3’UTR ARE were enriched for “mesenchyme development”, “cell migration” and “cell morphogenesis involved in differentiation” processes, among others (Suppl. Fig. 3C and gene list in Suppl. Table S1). In addition to the above rG4 and ARE motifs, we noticed that C/U-rich motifs were highly ranked in both 5’ and 3’UTRs but the sequences were not exactly identical (Fig. 4J). Nevertheless, by RNAshapes [34], these sequences were predicted to form very similar secondary structure featured by continuous hairpins (Hs) with flanking stems (Ss) (Suppl. Fig. 3D-E), suggesting that Dhx36 may recognize other specific secondary structures besides G4 and ARE. Lastly, on the binding regions of CDS, 8 nt binding motifs were also identified (Suppl. Fig. 3F) which interestingly had similarity with the binding motif for splicing regulators including SRSF1, SRSF2, TRA2A, TRA2B and HNRNPA2B1 (Suppl. Fig. 3G), suggesting a possibility for Dhx36 involvement in splicing regulation. Similarly, we also noticed that Dhx36 binding sites in introns (Suppl. Fig. 3H and Suppl. Table S1) were enriched in a motif resembling the C box motif (RUGAUGA) of C/D box snoRNAs (Suppl. Fig. 3I) and some of the binding sites were found overlapping with either annotated (52/234) or predicted snoRNAs by snoSCAN [35](21/182), indicating an intriguing connection between Dhx36 and snoRNAs, which was previously reported for other DEAD or DEAH box RNA helicases [36]. Altogether, the above global profiling of Dhx36 binding to mRNA has provided an unprecedented view of endogenous Dhx36 binding in myoblasts, demonstrating the predominant binding of Dhx36 to rG4 structures in 5’UTRs of transcripts meanwhile revealing previously unknown aspects of Dhx36 involvement in mRNA metabolism.

### Translational profiling reveals that Dhx36 facilitates mRNA translation through binding to 5’UTR rG4

The enrichment of Dhx36 binding at rG4 sites of 5’UTR suggested its potential involvement in global translational regulation by unfolding the 5’UTR rG4 in myoblasts. Accordingly, we performed polysome profiling coupled with RNA-seq to examine the translational status of mRNAs in C2C12 myoblasts with *Dhx36* deleted by CRISPR/Cas9 (Dhx36 KO) (Fig. 5A). Briefly, Dhx36 KO or Ctrl cell lysates were sedimented into supernatant, small subunits (40S), large subunits (60S), monosomes (80S) and polysomes (associated with ≥5 ribosomes thus considered to undergo active translation) by sucrose density centrifugation and fractionation (Fig. 5A). As shown in Figure 5B, the global absorbance profiles of ribosomes showed no obvious difference between Ctrl and KO cells, suggesting that Dhx36 does not affect global protein synthesis, which is consistent with a prior report in HEK293T cells in which knock-down of *Dhx36* did not have a major impact on polysome absorbance profiles [37].

**Figure 5.**
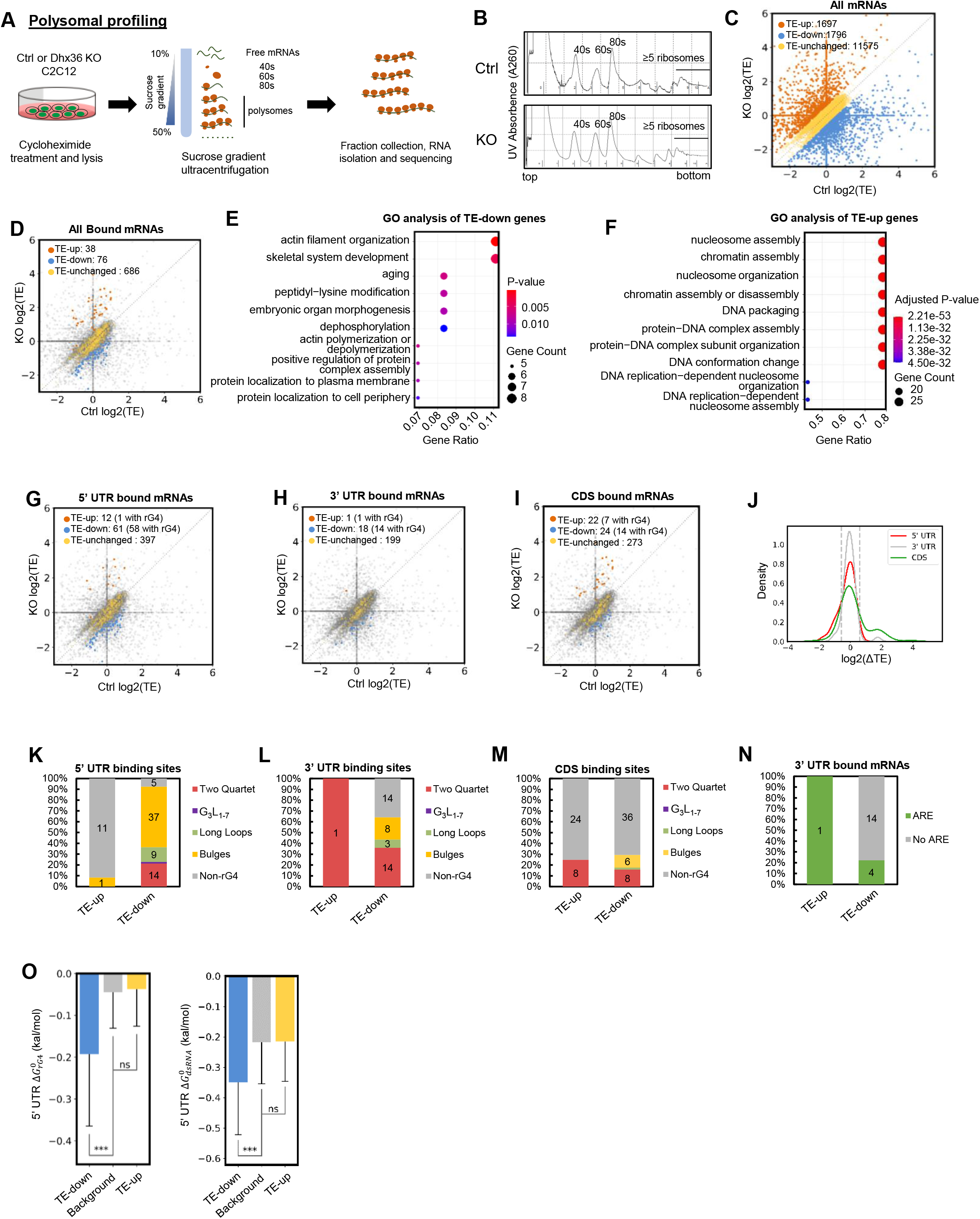
Translational profiling reveals that Dhx36 facilitates mRNA translation through binding to 5’UTR rG4. (A) Workflow for Polysome profiling performed in Ctrl or *Dhx36* KO C2C12 myoblast cells. (B) The Polysome profiles in Ctrl or KO reveal no global changes of translation by *Dhx36* loss. 10%-50% sucrose gradient was applied to achieve separation of polysomes. The peaks for small (40S) and large (60S) subunits, as well as monosome (80S), ≥5 ribosomes are indicated. (C) The translational efficiency (TE) in Ctrl or KO was calculated and the scatter plot of log2(TE) values of all mRNAs (n=15068 with consistent changes in two biological replicates) is shown. The numbers of mRNAs with up- (≥ 1.5 fold in KO vs Ctrl, orange dots) or down-regulated (≤ 0.66 fold in KO vs Ctrl, blue dots) or unchanged (yellow) TE are shown. (D) The above analysis was performed on all Dhx36 bound mRNAs. (E-F) GO analysis result for TE-up or - down genes in Figure 5D. (G-I) Scatter plot showing log2(TE) in KO vs Ctrl in 5’UTR, 3’UTR or CDS bound transcripts. (J) Kernel Density Estimates on log2(ΔTE) values of mRNAs with Dhx36 5’UTR, 3’UTR or CDS only binding. ΔTE: the TE alteration in KO vs Ctrl. (K-M) Prediction of rG4 formation in binding sites on 5’UTR, 3’UTR, CDS bound mRNAs was conducted and the number of binding sites possessing each subtype is shown in TE-up or ‒down mRNAs. (N) Prediction of ARE sites in binding sites on 3’UTR bound mRNAs was conducted and the number of mRNAs possessing ARE is shown. (O) 5’ UTR region predicted folding energies of rG4 and dsRNA structures in TE-up and TE-down genes with 5’UTR binding. One-tailed Mann-Whitney test was used to calculate the statistical significance between folding energy values: n.s., not significant (p>=0.05), ***P<0.001.

To further identify mRNAs whose translational efficiency (TE) might be affected by *Dhx36* loss, total mRNAs from the cell lysates and the fraction of mRNAs associated with polysomes were then collected for high-throughput sequencing. TE for each mRNA was calculated by normalizing the polysome-associated mRNA to total transcript level (see Materials and Methods). The log2 values of TE change in KO vs Ctrl (log2(ΔTE)) in the two biological replicates (Suppl. Fig. 4A and Suppl. Table S2, n=19179) were highly reproducible since large proportion of the mRNAs exhibited a similar TE change trend being up- or ‒down-regulated in both replicates (Supp. Fig. 4A, red dots, n=15068), with a Spearman Correlation Coefficient (SCC) of 0.956. Overall, among these 15068 mRNAs, 1697 displayed up-regulated (>=1.5 fold) (TE-up) TE values upon *Dhx36* loss while 1796 down-regulated (<=0.66 fold) (TE-down) (Fig. 5C and Suppl. Table S2). GO analysis revealed that the TE-up genes were enriched in processes such as “protein-DNA complex assembly” or “chromatin assembly”, and the TE-down genes were enriched in “pattern specification”, “positive regulation of neurogenesis”, among others. (Suppl. Fig. 4B-C). To further elucidate the TE changes caused by direct Dhx36 binding, the above data were intercepted with Dhx36 binding targets from the CLIP-seq, uncovering a much higher number (76 vs 38) of TE-down (<=0.66 fold) vs TE-up (>=1.5 fold) transcripts with Dhx36 binding (Fig. 5D and Suppl. Table S2), thus suggesting a promoting effect of Dhx36 on global mRNA translation. GO analysis showed that the TE-down genes were enriched in “skeletal system development” and “aging”, while the TE-up genes were enriched in “nucleosome assembly” and “chromatin assembly”, among other processes (Fig. 5E-F). Supporting that this new translational effect is largely related to 5’UTR binding, we found that Dhx36 binding to 5’UTRs conferred a stronger effect on TE change (a total of 73 transcripts with TE changes, Fig. 5G) as compared to bindings within 3’UTRs (19) and CDS (46) (Fig. 5H-I). Furthermore, among the 73 transcripts, 61 showed TE-down while 12 TE-up (Suppl. Table S2), suggesting that Dhx36 binding on 5’UTR principally promotes translational efficiency. On the other hand, Dhx36 binding within 3’UTR may also increase TE as 1 TE-up vs 18 TE-down mRNAs were found upon *Dhx36* loss (Fig. 5H and Suppl. Table S2). Interestingly, Dhx36 binding within CDS regions appeared to confer both increase and decrease of translation as about an equal number (22 vs 24) of transcripts were up-or down-regulated upon *Dhx36* loss (Fig. 5I and Suppl. Table S2). In line with the above findings, analyses with Kernel Density Estimate showed slightly lower log2(ΔTE) values were associated with 5’UTR bound mRNAs compared with values of 3’UTR or CDS bound ones (Fig. 5J).

To further confirm that the TE-promoting effect by Dhx36 is conferred by 5’UTR rG4 structures, we found indeed that a large number (58/61) of TE-down mRNAs with 5’UTR Dhx36 binding contained rG4 structures; a closer look revealed that the majority (61/66) of these 5’UTR binding sites possessed bulges as the dominant rG4 structure (37) (Fig. 5K); on the contrary, only one TE-up mRNA with 5’UTR Dhx36 binding contained rG4 structure. When the above analysis was performed on 3’UTR bound mRNAs, some correlation was found between rG4 formation and the number of TE-down transcripts (14/18) (Fig. 5H), which was even weaker on CDS (14/24) (Fig. 5I). Furthermore, the rG4 structures appeared again dominated by bulges and two quartets (Fig. 5L-M). In addition, the TE changes in 3’UTR bound genes had no clear correlation with the presence of ARE (only 5/19 mRNAs contained AREs) (Fig. 5N). Furthermore, we found much lower than background rG4 folding energy, 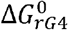 (i.e., more stable folded rG4 structures) in the TE-down but not in the TE-up mRNAs with 5’UTR binding. In addition, the sequence context favors the formation of rG4 structures over dsRNAs as 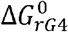 displayed a much larger fold difference compared with background (mean energy value 4.3 fold, p=2.64e-23) than 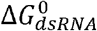 (mean energy value 1.61 fold, p=1.365e-16) (Fig. 5O). Taken together, these findings revealed that rG4 structures in 5’UTRs are a key determinant for the promoting function of Dhx36 in mRNA translation.

### *Dhx36* loss exerts a potential effect on total mRNA abundance

To investigate whether Dhx36 could modulate mRNA abundance, we next explored the global effect of *Dhx36* loss on total mRNA abundance using the 15068 genes with similar TE change trend (Suppl. Fig. 4A, red dots and Suppl. Table S3). Only a small number of mRNAs were up- (125, fold>=1.5) and down-(121, fold<=0.66) regulated in *Dhx36* KO vs Ctrl cells (Fig. 6A and Suppl. Table S3). The mRNA-up genes were enriched for GO terms including “tube morphogenesis”, “regulation of cytoskeleton organization” while mRNA-down genes were associated with processes such as “nucleosome assembly” and “chromatin assembly” (Suppl. Fig. 5A-B). Interestingly, binding within CDS and 3’UTR confers a stronger effect on mRNA abundance than binding within 5’UTR as a total of 99, 92 vs 63 mRNAs were altered, respectively (Fig. 6B-D, Suppl. Table S3). Specifically, 47 5’UTR bound mRNAs were up-regulated while 16 down-regulated, suggesting that 5’UTR binding decreases mRNA abundance. For both 3’UTR (37 up vs 55 down) and CDS (36 up vs 63 down) bound mRNAs, the trend was nevertheless opposite, suggesting that Dhx36 binding on 3’UTRs and CDS may likely cause increase in mRNA abundance. Similar conclusions can be drawn when further plotting the Kernel Density Estimate of the log2(ΔFPKM-total) values from transcripts with only 5’UTR, 3’UTR or CDS bindings (Fig. 6E). When further examining the correlation between rG4 formation with the abundance change of mRNAs, we found a particularly strong correlation between rG4 in 5’UTR with mRNA increase upon *Dhx36* loss (43/47 up-regulated transcripts possessed rG4s, Fig. 6B), implying that Dhx36 binding to 5’UTR rG4 may lead to decreased mRNA level. Although Dhx36 binding to 3’UTR and CDS appeared to increase mRNA levels, it is hard to conclude whether this is connected to rG4 binding as only 27/55 and 29/63 transcripts contained rG4. In all the rG4s residing in the described binding sites, bulges and two quartets, but not canonical rG4 motifs dominated (Fig. 6F-H). Similar to Fig. 5O, when computing the 5’UTR 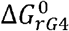 and 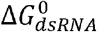, we found much lower than background 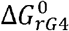 from the mRNA-up genes (mean energy value 3.19 fold, p=4.09e-15 compared to 1.43 fold, p=4.11e-10), favoring the formation of rG4 structures than dsRNAs on 5’UTRs of these transcripts (Fig. 6I). We next investigated the potential connection between 3’UTR ARE presence and RNA abundance as Dhx36 binding to ARE is known to affect RNA stability[12]. Unexpectedly, no strong correlation was found between the presence of ARE and mRNA abundance in the 23/92 3’UTR binding transcripts that possessed AREs in the binding sites (Fig. 6J). However, by further computing the 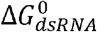 of 3’UTR AREs from Fig. 6J, AREs in both up- and down-mRNAs exhibited significantly lower than background folding energy values (Fig. 6K); the ambiguity of 3’UTR ARE in regulating mRNA abundance therefore deserves further investigation.

**Figure 6.**
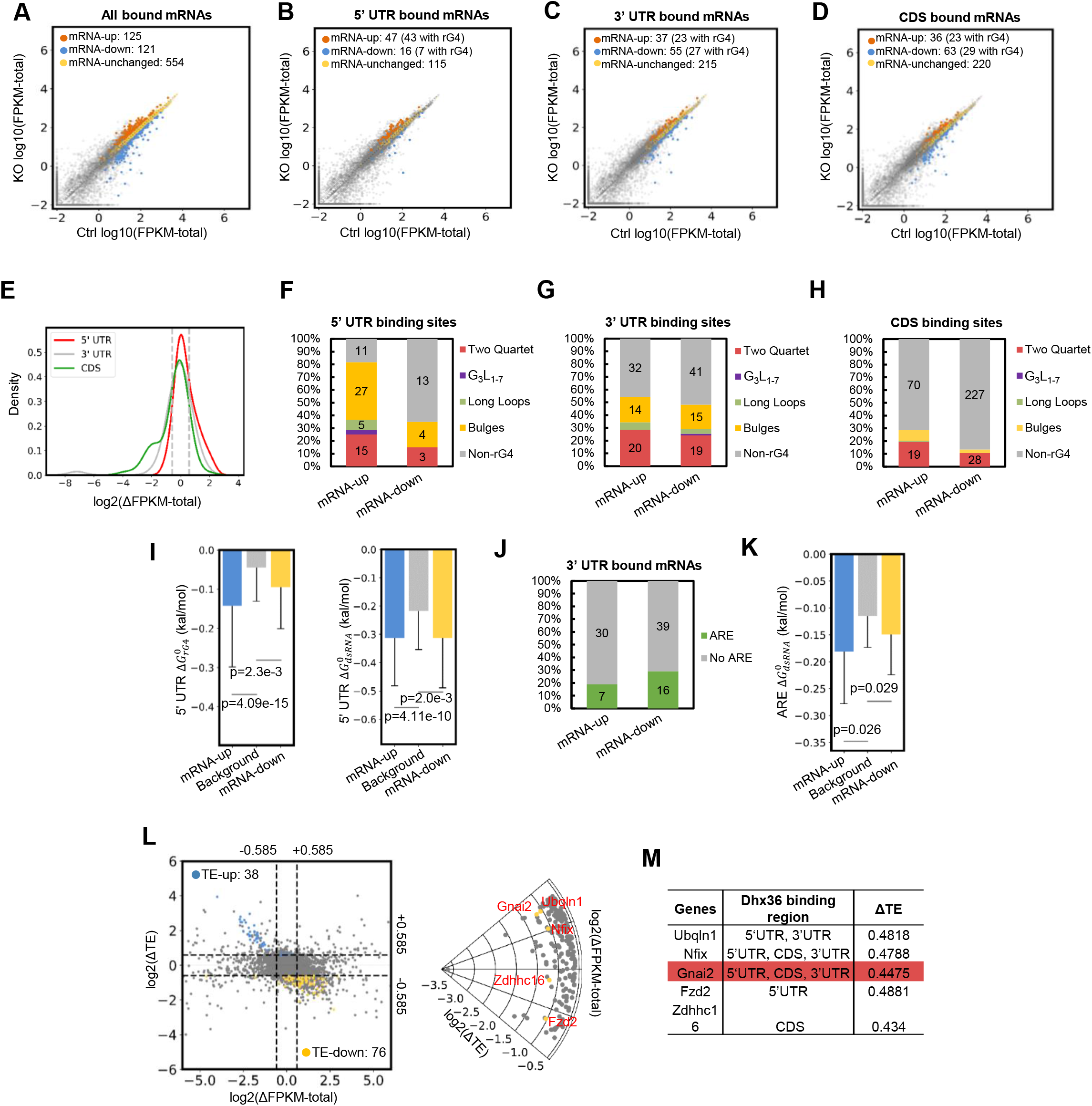
*Dhx36* loss exerts a potential effect on total mRNA abundance. (A-D) Scatterplot showing the total mRNA abundance (log10(FPKM-total)) values of all, 5’UTR, 3’UTR or CDS Dhx36 bound mRNAs in Ctrl vs KO; the number of up- (KO vs Ctrl ≥1.5 fold), down-(≤0.66 fold) or unchanged mRNAs are shown. (E) Kernel Density Estimates showing log2(ΔFPKM-total) values of mRNAs with only Dhx36 5’UTR, 3’UTR or CDS binding. ΔFPKM-total: the mRNA level alteration in KO vs Ctrl. (F-H) Prediction of rG4 formation in the binding sites of 5’UTR, 3’UTR, CDS bound mRNAs was conducted and the number of sites possessing each subtype is shown in up- or down- regulated mRNAs. (I) 5’ UTR region predicted folding energies of rG4 and dsRNA structures in up and down regulated mRNAs with 5’UTR binding. (J) The number of 3’UTR bound mRNAs possessing ARE sites is shown. (K) Predicted folding energies of dsRNAs in up- or down- regulated mRNAs with 3’UTR ARE binding. One-tailed Mann-Whitney test was used to calculate the statistical significance between folding energy values. (L) Left: Scatterplot illustrating TE (y-axis) and mRNA(x-axis) alterations in KO vs Ctrl with a threshold of ±0.585 (dashed line) for 7300 genes with FPKM-total values larger than 1 in either Ctrl or KO condition. Blue dots: TE-up genes with Dhx36 binding; yellow dots: TE-down genes with Dhx36 binding. Gray dots: the genes with no significant TE change or no Dhx36 binding. Right: A total of 5 bound genes (yellow dots) including *Ubq1n1, Nfix, Gnai2, Fzd2* and *Zdhhc16* (red font) showed constant mRNA and decreased TE levels in KO vs Ctrl using a more stringent threshold (log2(ΔTE) <−1, ΔTE<0.5). Gray dots: 197 genes with significant TE decrease and constant mRNA levels but with no Dhx36 binding. (M) Dhx36 binding regions and ΔTE values are shown for the 5 genes in Figure 6L and *Gnai2* is highlighted.

Lastly, to exclude the effect of mRNA change on TE decrease upon *Dhx36* loss, we compared the divergence at transcriptional and translation levels. As a result, among 114 Dhx36 bound genes with significant TE changes, 5 out of 38 TE-up (blue dots) and 16 out of 76 TE-down (yellow dots) genes showed constant mRNA level in KO vs Ctrl cells (Fig. 6L, left). By a stringent threshold of TE change <0.5, only 5 mRNAs, *Ubq1n1, Nfix, Gnai2, Fzd2 and Zdhhc16* were significantly decreased in TE upon *Dhx36* loss (Fig. 6M), suggesting that they are canonical translational regulatory targets of Dhx36 in myoblasts that merit further investigation.

### Dhx36 controls SC proliferation through regulating *Gnai2* translation by binding and resolving the 5’UTR rG4 site

Once our system-wide analyses uncovered Dhx36 as a major translation regulator through binding to 5’UTR rG4, we next aimed at elucidating the downstream effectors of its pro-proliferative function in myoblast by selecting a target transcript as a proof-of-principle (Fig. 6L, right). We chose *Gnai2* mRNA as it shows clear Dhx36 binding on the rG4 sites within its 5’UTR (Fig. 6M, Suppl. Fig. 6A and Fig. 7A-B). Of note, previous studies showed that constitutive activation of Gnai2 resulted in muscle hypertrophy and accelerated injury-induced muscle regeneration while knocking-out *Gnai2* reduced muscle size and impaired muscle regeneration [38, 39], which to some degree phenocopies the impaired muscle regeneration in *Dhx36* mutant mice, suggesting that *Gnai2* is a likely target of Dhx36.

**Figure 7.**
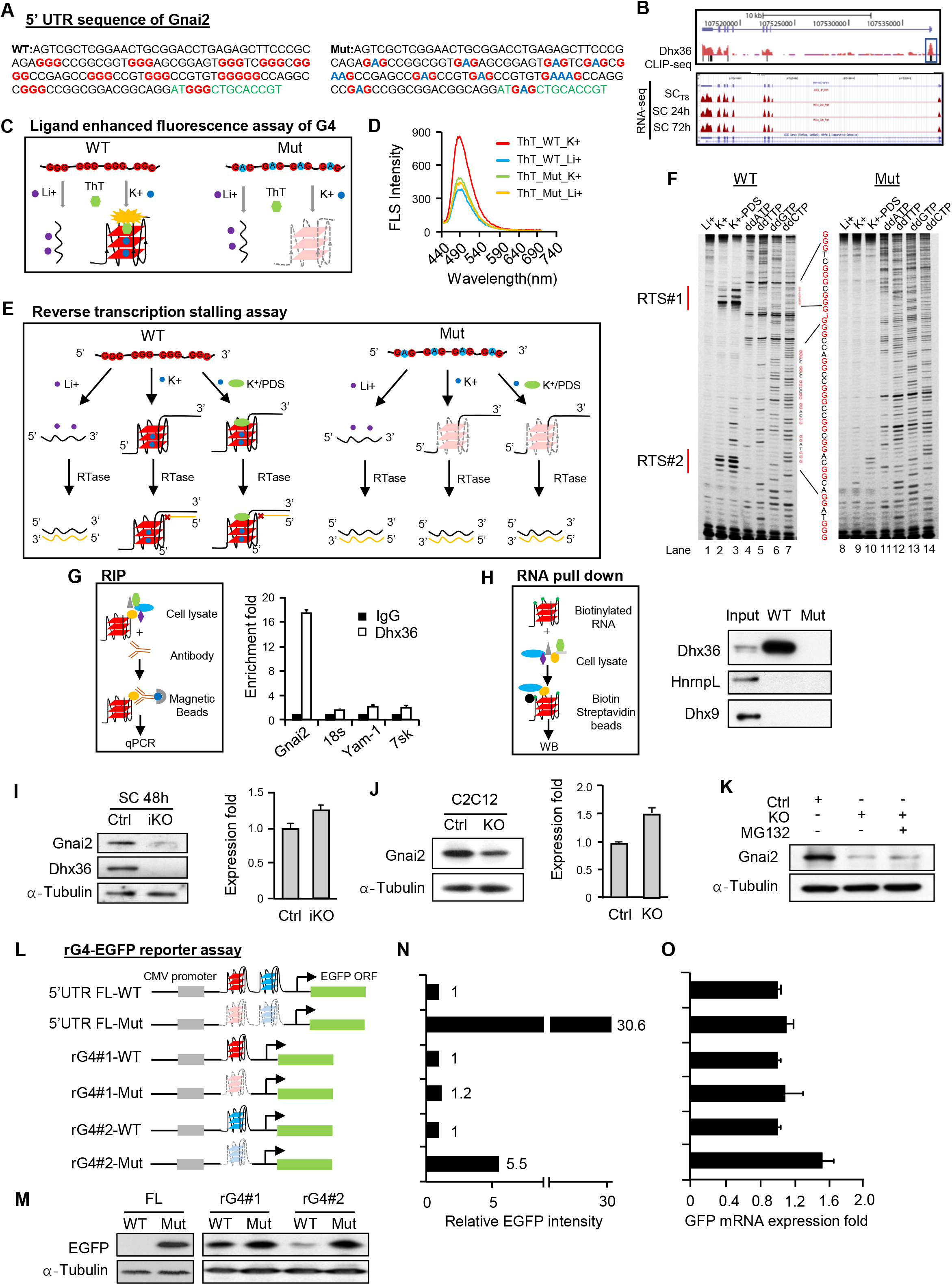
Dhx36 regulates *Gnai2* mRNA translation by unwinding the rG4 formed at the 5’UTR. (A) Sequence of the 5’UTR of WT or mutant (Mut) *Gnai2* mRNA. Green color denotes coding sequence. All GGGs are highlighted in red. Mutated Gs are highlighted in blue. (B) Top: genomic snapshot of Dhx36 CLIP-seq track showing its binding at *Gnai2* mRNA. Bottom: snapshot of RNA-seq tracks showing *Gnai2* mRNA levels in freshly isolated SCs, SCs cultured for 24h or 72h. Illustration of the Thioflavin T (ThT) staining assay. WT or Mut *Gnai2* 5’UTR RNA were treated with 150 mM Li+ or K+ together with 1 μM ThT and excited at 425nm. (D) Plot of the intensity of Fluorescent spectrum (FLS) with the wavelength from 440nm to 700nm. (E) Schematic illustration of Reverse transcriptase stalling (RTS) assay. WT and Mut *Gnai2* 5’UTR RNAs were treated with 150 mM Li+, 150 mM K+ or 150 mM K+ plus 2 μM PDS. Reverse transcriptase was stalled at the rG4 formation site to form reverse stalling site (RTS). (F) The cDNAs generated by the above reverse transcription were run on a Dideoxy sequencing gel. Left: In WT *Gnai2* 5’UTR RNA, two major RTS (highlighted as RTS #1 and #2) sites were identified. Right: no RTSs were detected in Mut *Gnai2* 5’UTR RNA. (G) RNA immunoprecipitation (RIP) (illustrated in the left panel) was performed with Dhx36 antibody and the retrieved *Gnai2* transcripts were detected by qRT-PCR. *18s, Yam-1* and *7sk* transcripts were used as negative controls. (H) RNA pull-down (illustrated in the left panel) was performed with *in vitro* transcribed and biotin-labeled WT or Mut *Gnai2* 5’UTR RNAs and the retrieved Dhx36 protein was detected by Western blot. HnrnpL and Dhx9 were used as negative controls. (I) Left: Gnai2 protein was detected by Western blot in Ctrl and iKO SCs cultured for 48h. α-Tubulin was used as internal loading control. Right: *Gnai2* mRNA was detected by qRT-PCR in the above cells. *18s rRNA* was used as normalization control. (J) Gnai2 protein and mRNA were detected in proliferating Ctrl and Dhx36 KO C2C12 cells. (K) Gnai2 protein was detected in Ctrl, KO or KO C2C12 cells treated with proteasome inhibitor MG132 for 8 h. (L) Cloning scheme of the plasmids used in the reporter assay. WT or Mut sequences of full length 5’UTR, rG4#1, or rG4#2 only were cloned upstream of EGFP open reading frame in pEGFP-N1 vector. All GGGs were mutated to GAG in the Mut sequences. (M) The above plasmids were transfected into C2C12 cells and EGFP protein was detected 48 h post transfection by Western blot. A-tubulin was used as the internal loading control. (N) The relative intensity of each EGFP protein band normalized by α-Tubulin band was calculated. The intensity for the WT was set as 1. (O) EGFP mRNA was detected by qRT-PCR from the above samples.

To validate the rG4 formation in the identified site of *Gnai2* 5’UTR, we found that the G-rich region is very conserved among multiple species including Mouse, Human, Chimp, Rhesus Monkey and Cow (Suppl. Fig. 7A). By probing with Thioflavin T (ThT), which becomes fluorescent in the presence of the G-quadruplexes [40], the 5’UTR of WT *Gnai2* exhibited an enhanced emission at the wavelength of 490nm in K+ (which is known to stabilize G4 structure) compared to in Li+; as expected, when all GGGs in the site were mutated to GAGs (Fig. 7 Mut) and the formation of G-quadruplexes was disrupted, the enhanced emission was largely lost in K+ (Fig. 7C-D). Moreover, probing with another G4 ligand *N*-methylmesoporphyrin (NMM) [41, 42] also showed that the WT but not Mut 5’UTR could form rG4 in K+ (Suppl. Fig. 7B). Lastly, targeted G4 formation of *Gnai2* 5’UTR was also examined by RTS (reverse transcriptase stalling) gel analysis [43] which used WT or mutated RNA as the templates for reverse transcription at Li+, K+ or K+ plus PDS (pyridostatin, a G4 ligand which can further stabilize G4 structure) conditions to detect the stalling caused by the formation of G4 structures (Fig. 7E). Two very strong stalling sites (RTS#1 and RTS#2) were detected for WT 5’UTR under G4 formation conditions but not the mutated RNA (Fig. 7F). Altogether, the above results confirmed the formation of rG4 structure at *Gnai2* 5’UTR.

Next, to validate the interaction between the above identified rG4 site and Dhx36, we performed RNA immunoprecipitation experiment and showed that the Dhx36 antibody effectively retrieved *Gnai2* mRNA but not control RNAs including *18s, Yam-1 or 7sk* (Fig. 7G). Consistently, RNA pull-down assay using WT 5’UTR sequence retrieved a significant amount of Dhx36 protein, which was completely lost when GGG was mutated to GAG, strengthening that Dhx36 does interact with *Gnai2* 5’UTR through the rG4 site (Fig. 7H).

To further investigate the translational control exerted by Dhx36 in myoblasts, we found that Gnai2 protein but not mRNA was significantly reduced in both activated SCs from iKO mice compared with those from Ctrl (Fig. 7I) and proliferating KO C2C12 compared with control myoblast cells (Fig. 7J). Furthermore, when the *Dhx36* KO cells were treated with MG132, the inhibitor of proteasome, Gnai2 protein level was not increased (Fig. 7K), suggesting that protein degradation of Gnai2 was not affected by *Dhx36* loss. To further study whether Dhx36 modulation of *Gnai2* mRNA translation is mediated by resolving rG4 formation, we generated a GFP reporter by inserting the WT or mutated full-length 5’UTR of *Gnai2,* rG4#1 or rG4#2 sequence to the upstream of the GFP ORF in a pEGFP-N1 plasmid [19] (Fig. 7L). When transfected into C2C12 myoblasts, the protein levels of GFP of all three WT reporters were drastically decreased compared with the correspondent mutated reporters as measured by Western blot for EGFP protein (Fig. 7M) with the relative intensity of EGFP band calculated by ImageJ (Fig. 7N), while the level of GFP mRNA was not significantly altered (Fig. 7O), indicating that the rG4 formation in *Gnai2* 5’UTR indeed regulates mRNA translation.

Lastly, to test the functional connection of Dhx36 and Gnai2 in SC proliferation, we found that transduction of SCs with Gnai2 expressing lentivirus promoted cell proliferation as assessed by a 26.9% increase in EdU+ cells (Fig. 8A-B). Conversely, when SCs were transfected with siRNAs against *Gnai2*, a modest but significantly decreased level of EdU incorporation was observed (Fig. 8C-D). The above results thus suggested that Dhx36 and Gnai2 conform a regulatory axis for proliferation of SCs. In fact, transduction of SCs from *Dhx36* iKO with a Gnai2-expressing virus led to a 28.5% increase in EdU+ cells compared to SCs infected with Ctrl virus (Fig. 8E-F), suggesting that Gnai2 is indeed a downstream effector of Dhx36 in promoting SC proliferation.

**Figure 8.**
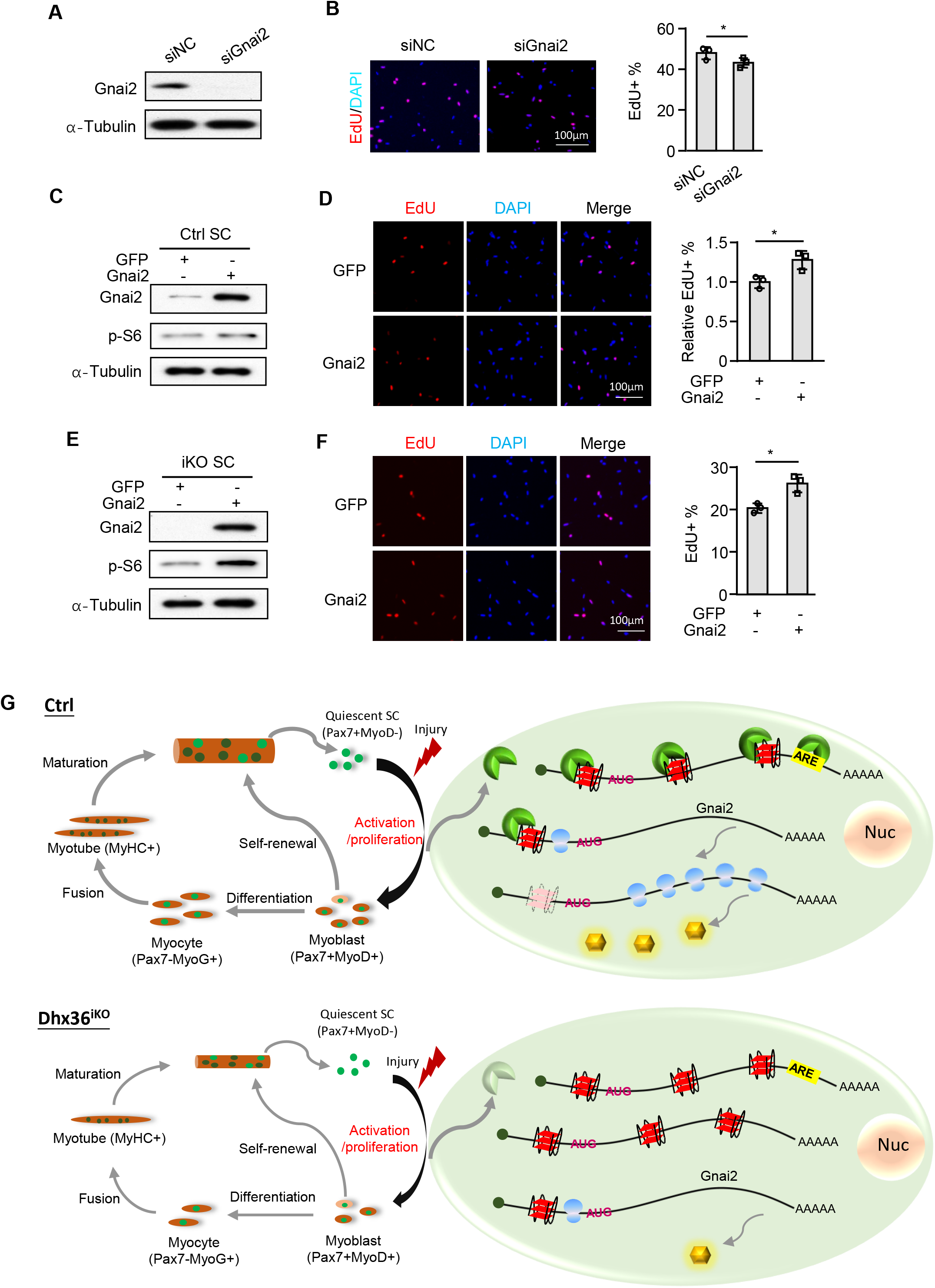
Gnai2 mediates Dhx36 function in SC proliferation. (A) C2C12 cells were transfected with siNC or siGnai2 oligos and harvested at 48 h post-transfection. The successful deletion of Gnai2 protein was detected by Western blot with α-Tubulin as the internal loading control. (B) SCs were transfected with the above siRNA oligos for 48 h when EdU were added to the culture medium 4 h before staining. The percentage of EdU incorporation was quantified from 10 randomly selected fields/sample. Scale bar=100 μm; n=3 per group. (C) Ctrl SCs were infected with GFP or Gnai2 expressing virus and the Gnai2 overexpression was confirmed by Western blot at 48 h after infection. EdU was added to the above cells for 4 hours and the relative EdU incorporation percentage was quantified from 10 randomly selected fields/sample with the value from GFP virus infected cells was set as 1. Representative images are shown. Scale bar=100 μm; n=3 mice per group. (E) iKO SCs were infected with GFP or Gnai2 expressing virus and the expression of Gnai2 protein was confirmed by Western blot at 48h after infection. (F) EdU labeling was performed as above described and the incorporation percentage was quantified from 10 randomly selected fields/sample. Representative images are shown. Scale bar=100 μm; n=3 mice per group. Data represent the average of three independent experiments□±□s.d. Student’s t-test (two-tailed unpaired) was used to calculate the statistical significance in N, Student’s t-test (two-tailed paired) was used to calculate the statistical significance in P and R: *, P < 0.05. (G) Schematic illustration of the functional mechanism of Dhx36 in regulating SC proliferation and muscle regeneration.

## Discussion

In this study, we demonstrate a fundamental role of Dhx36, an RNA G-quadruplex (rG4) structure unwinding helicase, in skeletal muscle stem cells during muscle regeneration, via post-transcriptional regulation of mRNA metabolism. We found that specific inactivation of Dhx36 in SCs impaired muscle regeneration through inhibiting their proliferative expansion. By combined CLIP-seq and polysome profiling, we discovered that Dhx36 facilitates mRNA translation by binding and unwinding the rG4 structures formed at the 5’UTR regions of target mRNAs. In particular, Gnai2 is a direct downstream effector of Dhx36 proliferation-promoting functions. Beyond translational control, our analyses also reveal potential functions of Dhx36 in regulating mRNA abundance and other previously unknown aspects of mRNA metabolism. Our findings, in sum, uncover Dhx36 as a versatile RBP and highlight the importance of post-transcriptional regulatory mechanisms in stem cells and tissue regeneration.

While epigenetic and transcriptional control of myogenesis and muscle regeneration has been studied extensively, the importance of post-transcriptional regulation in these processes is much less defined. Post-transcriptional regulation of gene expression permits cells to orchestrate rapid changes in protein levels from steady-state mRNAs, thus ensuring that a pool of primed stem cells is available to repair and maintain tissue function. Recent evidence supports the importance of post-transcriptional regulation in SCs. For example, it is shown that some mRNAs are known to be transcribed and stored in the quiescent SCs to allow the rapid production of protein products once SC activated [9–11, 44]. Our findings for the first time provide solid genetic evidence demonstrating that Dhx36 is an indispensable post-transcriptional regulator of skeletal muscle regeneration. Despite the increasing number of rG4 interacting RBPs discovered to date [45–48], loss of *Dhx36* completely blunted muscle regeneration with no compensation by other helicases; this was confirmed by both constitutive and inducible loss of *Dhx36* in SCs *in vivo*. *Dhx36* loss caused a major defect in late SC activation and proliferation stages, which resulted in a drastic decrease in the cycling cell progeny required for muscle regeneration; this function is consistent with its expression dynamics, with *Dhx36* being induced as SCs become fully activated and proliferate (Fig. 1B-D). Prior studies also showed the involvement of Dhx36 in cellular proliferation, for example in erythrocytes [19, 22, 27, 49]. Nevertheless, we cannot exclude that Dhx36 may also regulate additional SC myogenic stages.

To fathom the mechanisms underlying Dhx36 function in SC biology, we integrated CLIP-seq binding and polysome profiling analyses. During the preparation of this manuscript, Sauer et al. published the very first transcriptome-wide profiling of Dhx36 in HEK293 cells by PAR-CLIP using FLAG or HA tagged plasmids overexpressing DHX36. As the dose of RBP may significantly modulate the RNA structure, and thus binding dynamics, obtaining the endogenous Dhx36 binding profile (our study) is critically needed to understand the physiological role of this helicase. Consistent with its almost excusive cytoplasmic localization (Fig. 1E), it is not surprising the majority of Dhx36 interacting transcripts are polyadenylated mRNAs while a small portion arise from intergenic regions. Expectedly, the highest binding density was located in the 5’UTR regions although the highest number of binding sites was found in CDS and introns (Fig. 4E-F). We demonstrated that the dominant binding motifs in 5’UTR, 3’UTR and CDS are all G-rich and predicted to form rG4s, strongly suggesting that Dhx36 function is highly dependent on rG4s at least in myoblasts. Interestingly, bulges, two quartet but not canonical rG4 structures were predicted, which suggests cell-type specific binding preference or dynamics for rG4 subtypes. Unlike Sauer’s study suggesting wild type DHX36 cannot bind to AU-rich sequences while the mutant form, with an inactive helicase (DHX36-E335A), is capable of, we found that endogenous Dhx36 in myoblasts interacts with 3’UTR AREs in a large set of transcripts. When integrated with polysome profiling, we were able to demonstrate that 5’UTR binding of Dhx36 indeed confers a promoting function in translational efficiency as a much higher number of Dhx36 bound genes were down-rather than up-regulated. Dhx36 binding to the 3’UTR and CDS has not been previously documented. Notably, our data on the Dhx36 binding with rG4 in the 3’UTR suggest a role on translation. Interestingly, Dhx36 binding on CDS had a marked effect on TE of a number of transcripts; this effect may be rG4 independent despite the knowledge that rG4 formation in CDS could impact translational elongation [50–52]. Altogether, our findings uncover a wide range of Dhx36 binding locations that may mediate diversified functional mRNA regulatory mechanisms either rG4 dependent or independent, which in turn impact stem cell biology.

In addition to the translational effect, we took efforts in exploring the Dhx36 possible impact on mRNA abundance. When trying to connect rG4 binding with the mRNA abundance regulation, we found a high percentage of up-regulated mRNAs possessing 5’UTR binding on rG4 sites, suggesting that *Dhx36* loss increases transcript abundance, which could result from retarded translation. Intriguingly, we did not find any connection between mRNA abundance and Dhx36 binding to 3’UTR ARE, despite that Dhx36 was first defined as a ARE bound RBP that regulates mRNA degradation[12]; Sauer et al. also failed to tease out whether ARE contributed to mRNA abundance changes[22]. The ambiguity of ARE in mediating Dhx36 function thus needs to be further investigated. Additionally, it will be worth investigating the rG4-dependent regulation of mRNA abundance by Dhx36. As rG4 could contribute to post-transcriptional processing and RNA decay, it is highly possible that Dhx36 has diversified functional mechanisms through its connection with rG4.

Lastly, we proved that Gnai2 is a downstream effector of Dhx36 in promoting SC proliferation. The mechanistic elucidation of the Dhx36-Gnai2 regulatory axis also served as a proof-of-concept for dissecting how Dhx36 binding to 5’UTR rG4 promotes translational initiation. Various biophysics and biochemical methods were employed to demonstrate the presence of the rG4 and its binding with Dhx36 as well as the regulation in translational initiation. We envision that many other targets, similar to Gnai2, exist in stem cells that mediate the pleiotropic Dhx36 functions, which calls for future intensive investigations.

## Materials and Methods

### Mice

*Pax7^CreER^* (*Pax7^tm1(cre/ERT2)Gaka^*)[53] and Tg: Pax7-nGFP mouse strains[23] were kindly provided by Dr. Shahragim Tajbakhsh. *Pax7^Cre^* (*Pax7^tm1(cre)Mrc^*) mouse was kindly provided by Dr. Charles Keller. *ROSA^EYFP^* mouse was provided by Jackson Laboratory. *Dhx36^fl/fl^* strain was kindly provided by Dr. Zhongzhou Yang with the authorization of Dr. Yoshikuni Nagamine who originally generated this mouse strain[19, 27]. *Pax7^Cre^* and *Dhx36^fl/fl^* mice were mated to generate *Dhx36* conditional KO (*Dhx36* cKO) mice (Ctrl: *Pax7^Cre/+^; Dhx36^+/+^*, cKO: *Pax7^Cre/+^; Dhx36^fl/fl^*). *Pax7^CreER^* mouse was crossed with *ROSA^EYFP^* mice to generate the *Pax7^CreER^; ROSA^EYFP^* reporter mice. The *Dhx36* inducible conditional KO (*Dhx36* iKO) mice with EYFP reporter (Ctrl: *Pax7^CreER/+^; ROSA^EYFP/+^; Dhx36^+/+^,* iKO: *Pax7^CreER/+^; ROSA^EYFP/+^; Dhx36^fl/fl^*) were generated by crossing *Pax7^CreER^; ROSA^EYFP^ with Dhx36^fl/fl^* mice. All animal handling procedures and protocols were approved by the Animal Ethics Committee at Chinese University of Hong Kong (CUHK).

### Animal procedures

Inducible conditional deletion of *Dhx36* was administered by injecting Tamoxifen (Tmx) (T5648, Sigma) intraperitoneally (IP) at 2mg per 20g body weight. For Bacl2 induced muscle injury, approximately 2 months old mice were intramuscularly injected with 50 μl of 10 μg/ml Bacl2 solution into tibialis anterior (TA) muscles and the muscles were harvested at designated time points for further analysis. For EdU incorporation assay *in vivo*, 2 days after Bacl2 injection, EdU injection via IP at 0.25 mg per 20g body weight was performed, followed by FACS isolation of SCs 12 hours later. Cells were then collected and fixed with 4% PFA. EdU labeled cells were visualized using “click” chemistry with an Alexa Fluor® 594 conjugated azide. Images were captured with a fluorescence microscope (Leica). For cardiotoxin (CTX) induced muscle regeneration, the tibialis anterior (TA), gastrocnemius and quadriceps muscles were injected with CTX (Latoxan; 10−5 M). At the indicated time points (3- and 7-day post injury), mice were euthanized, and muscles were either snap frozen or dissected for satellite cell isolation by FACS.

### Grip strength measurement

Grip strength of all four limbs was measured using a grip-strength meter (Columbus Instruments, Columbus, OH). The animal was held so that all four limbs paws grasped the specially designed mouse flat mesh assembly and the mouse was pulled back until their grip was broken. The force transducer retained the peak force reached when the animal’s grip was broken, and this was recorded from a digital display. Five successful grip strength measurements within 2 min were recorded. The maximum values were used for analysis. The mice were trained on the grip strength meter before the trial. Maximal muscle strength was obtained as values of KGF (kilogram-force) and represented in grams.

### Satellite cell sorting

Skeletal muscle tissue from hindlimb, forelimb and pectoralis muscles were dissected, gently minced with blades and digested with Collagenase II (1000U/ml) in Ham’s F10 for 90 minutes in a shaking water bath at 37°C. Digested tissue was then washed twice with rinsing media (10% HS, in Ham’s F-10) and centrifuged at 1500 rpm at 4°C for 5 minutes. Second digestion was performed by adding Collagenase II (1000U/ml) and Dispase (11U/ml) in rinsing media and incubated in shaking water bath at 37°C for 30 minutes. Digested tissue was passed through a 20-gauge needle for 12 times and filtered through a 40 μm filter followed by spinning at 1500 rpm for 5 min at 4°C. SCs were then sorted out by FACS Aria Fusion (BD) and collected as GFP+ groups. For isolating SCs without endogenous YFP expression, PE-Cy7-conjugated anti-CD31 (Biolegend, 102418), anti-CD11b (Biolegend, 101215/16) and anti-Sca-1 (Biolegend, 108113/14) antibodies were used to exclude the Lin (−) negative population. AF647-conjugated anti-CD34 (BD Pharmigen, 560230) and PE-conjugated anti-α7-integrin (Ablab, AB10STMW215) were used for double-positive staining of quiescent satellite cells or single α7-integrin staining of proliferating satellite cells. Isolated satellite cells were used either for RNA/protein extraction or were cultured in Ham’s F10 supplemented with 20% FBS and bFGF (0.025 μg/ml) (growth medium) for analysis of Dhx36 expression in quiescence, *in vitro* activation and proliferation.

### Single muscle fiber *ex vivo* culture

Single myofibers were isolated from the extensor digitorum longus (EDL) muscles of 2-3 months old mice by dissociation with collagenase II solution (800U/ml) at 37°C in a water bath for 75 min. Dissociated single myofibers were manually collected and transferred to a new dish and the operation was repeated for 2-3 times to remove dead fibers and debris. Isolated single myofibers were then either fixed in suspension immediately in 4% paraformaldehyde (PFA) for 15 min or maintained in suspension culture. For EdU incorporation assay, EdU was added to single myofibers which were cultured for 30 hrs and pulsed for another 4 hours. Fibers were then fixed in 4% PFA for 15 min and stained following the EdU staining protocols provided by manufacturer (Thermo Fisher Scientific, C10086).

### Cells

Mouse C2C12 myoblast cells (CRL-1772) and 293T cells were obtained from American Type Culture Collection (ATCC) and cultured in DMEM medium with 10% fetal bovine serum, 100 units/ml of penicillin and 100 μg of streptomycin (growth medium, or GM) at 37 °C in 5% CO2. For *in vitro* differentiation, C2C12 myoblasts were plated and cultured to high confluence and changed to the differentiation media (DM) (DMEM with 2% horse serum). Cells were cultured in DM and harvested for Western blot at day 1, 3 and 5. WT primary myoblast cells were cultured in Ham’s F10 medium supplemented with 20% FBS and bFGF (0.025 mgr/ml) (growth media, GM). For *in vitro* differentiation experiments, myoblasts were plated and cultured to high confluence and changed to DMEM with 5% Goat Serum (differentiation media, DM). Cells were cultured in DM during 24, 48, 72 and 96 h. For generating *Dhx36* knockout (KO) C2C12 cells, two sgRNAs expressing plasmids[54] which targeted the third exon of *Dhx36* were transfected to C2C12 and clones were selected by genotyping. Three homozygous C2C12 clones were selected for functional assays. Lenti viruses expressing GFP or Gnai2 were packaged in 293T cells as previously described. Ctrl or iKO SCs were infected by GFP or Gnai2 expression virus after attaching to the culture plate. 48h after infection cells were used for EdU incorporation assay, collected for Western blot or qRT-PCR.

### Preparation of DNA oligonucleotides

All DNA oligonucleotides were purchased from Integrated DNA Technologies (IDT). The Cy5 fluorescence-labelled DNAs were purified by HPLC. The quality of each oligonucleotide was confirmed by ESI-MS by IDT, with a single peak at the expected size, thus all DNA oligonucleotides were used without further purification. The sequences of the oligos can be found in Suppl. Table S4.

### Plasmids

Two sgRNAs for generating *Dhx36* knockout (KO) C2C12 cells were designed by CRISPOR[54] to target the third exon of *Dhx36* and cloned into PX458 vector at BbsI site. For lentivirus package and infection, mouse Gnai2 ORF were amplified from C2C12 cell cDNA and cloned into pLenti vector through PmeI and EcoRI restriction sites. For GFP reporter assays, the wild type full-length 5’UTR of Gnai2 or the mutated one with all the GGG mutated to GAG were cloned to the pEGFP-N1 vector at the XhoI/EcoRI sites. WT or Mut rG4#1 or rG4#2 sequenced were synthesized by BGI and cloned directly to pEGFP-N1 vector. The sequences of the cloning primers and oligos are shown in Suppl. Table S4.

### GFP reporter assay

The GFP reporter assay was conducted as previously reported[19]. Reporter vectors were transfected to the C2C12 cell lines and cultured for 48 hrs before harvested for RNA and protein extraction. Western blot and qRT-PCR was used to evaluate the protein and mRNA level of GFP respectively.

### Real-time PCR

Total RNAs from SCs or C2C12 cells were extracted using TRIzol reagent (Life Technologies) according to the manufacturer’s protocol. QRT-PCR was performed by using SYBR Green Master Mix (Applied Biosystem) on ABI PRISM 7900HT (Applied Biosystem). 18s rRNA or GAPDH mRNA was used for normalization. All the experiments were designed in triplicates. Primers used for qRT–PCR are shown in Supplementary Table S4.

### RNA pull-down assay

RNA pull down assay was conducted as previously described[55, 56]. Briefly, biotin-labeled RNAs were *in vitro* transcribed with the Biotin RNA Labeling Mix (Roche) and T7/T3 RNA *in vitro* transcription kit (Ambion) and purified with RNAeasy kit (Qiagen). C2C12 cells were harvested and lysed with RIPA buffer. Pre-folded RNA was then mixed with cell lysate and incubated at RT for 1 hour. Streptavidin agarose beads (Invitrogen) were then added to the binding mix and incubated for another hour. Beads were washed in Handee spin columns (Pierce) and binding proteins were retrieved from beads and used for Western blot analysis.

### RNA Immunoprecipitation

RNA Immunoprecipitation assay was conducted as previous description[55, 57]. 2μg of antibodies against Dhx36 (Abcam, ab70269) or isotype IgG (Santa Cruz Biotechnology) were used in this assay. Pulled-down RNA were resuspended in 20ul of RNase-free water and cDNAs were obtained from reverse transcription. QRT-PCR was performed with cDNAs by using SYBR Green Master Mix (Applied Biosystems). Relative enrichment was calculated as the amount of amplified DNA normalized to the values obtained from IgG immunoprecipitation which was set as 1.

### Preparation of *in vitro* transcribed RNAs

*In vitro* transcribed (IVT) RNAs were made using pre-annealed DNA hemiduplex and HiScribe™ T7 High Yield RNA Synthesis Kit following the manufacturer’s protocol. Each RNA was purified using the 7M urea, 15% denaturing acrylamide gel (Life Technologies) and the desired RNA gel band was sliced under UV. The gel piece was crushed and soaked in 1X 10 mM Tris pH 7.5, 1 mM EDTA, 800 mM LiCl (1X TEL800). The mixture was under constant shaking at 1300 rpm overnight at 4 °C. Next, the mixture was filtered against 0.22 μM filter and purified using RNA Clean & Concentrator-5 Kit (Zymo). The IVT RNA was then stored at −20 °C before use.

### Reverse transcriptase stalling assay

Reverse transcriptase stalling assay was performed similar to what was reported previously[43]. Three to five pmol of IVT RNA was added up to 4.5 μL with nuclease-free water, and 1 μL of 5 μM Cy5 fluorescence-labelled was added. The mixture was heated at 75 °C for 3 min, followed by incubation at 35 °C for 5 min. At the beginning of the 35 °C incubation, 3 μL of reverse transcription buffer was added to reach a final concentration of 150 mM KCl, 4 mM MgCl_2_, 20 mM Tris pH 7.5, 1 mM DTT, and 0.5 mM dNTPs. For cation-dependent experiments, either 150mM KCl or LiCl was used, unless otherwise stated. For ligand-dependent experiments, 1 μL of 20 μM PDS was added after the reverse transcription buffer. The 9.5 μL mixture was heated up to 55 °C and 0.5 μL of Superscript III (200U/ μL) was added to make up the 10 μL reaction. The reverse transcription mixture was incubated at 55 °C for 15 min, and then 0.5 μL of 2M NaOH was added at the end of the step. Then the mixture was incubated at 95 °C for 10 min to inactivate the SSIII and degrade the RNA template. For RTS, 10 μL of 2X stopping dye solution which contains 20 mM Tris, pH 7.5, 20mM EDTA, 94% deionized formamide was added to the reaction mixture. Orange G dye was added as tracker.

### Fluorescence spectroscopy

Fluorescence spectroscopy was performed similar to what was reported previously[58, 59]. Briefly, 1 μM RNAs in a reaction volume of 100 μL were mixed in 10 mM LiCac (pH 7.0) buffer and 150 mM KCl or LiCl. Samples were annealed at 95 °C for 5 minutes and cooled to room temperature for 15 minutes for renaturation. 5 μL of 20 μM NMM or ThT were added to the samples and samples were excited at 394 nm and 425 nm for NMM and ThT ligands respectively. The emission spectrum was collected from 550 – 750 nm for NMM and 440 nm – 700 nm for ThT. 1-cm path length quartz cuvette was used and measurements were performed using HORIBA FluoroMax-4. Spectra were acquired every 2 nm at 25 °C for WT and Mut samples. The entrance and exit slits were 5 and 2 nm, respectively.

### Immunoblotting and Immunofluorescence

For Western blot assays, *in vitro* cultured cells were harvested, washed with ice-cold PBS and lysed in cell lysis buffer. Whole cell lysates were subjected to SDS–PAGE and protein expression was visualized using an enhanced chemiluminescence detection system (GE Healthcare, Little Chalfont, UK) as described before[30]. The following dilutions were used for each antibody: Dhx36 (Abcam; 1:5,000), α-Tubulin (Sigma; 1:5000), MyoD (Santa Cruz Biotechnology; 1:1000), Gnai2 (Abcam; 1:5,000), CCND1 (Santa Cruz Biotechnology; 1:5000), CCNA1 (Santa Cruz Biotechnology), p-S6 (Santa Cruz Biotechnology; 1:5,000), HnrnpL (Santa Cruz Biotechnology; 1:5,000), Dhx9 (Santa Cruz Biotechnology; 1:5,000). For immunofluorescence staining, cultured cells and myofibers were fixed in 4% PFA for 15 minutes and blocked with 3% BSA within 1 hour. Primary antibodies were applied to samples with indicated dilution below and the samples were kept at 4°C overnight. For Dhx36 staining, FACS-sorted satellite cells were plated in 15-well chambers (IBIDI) and cultured in Ham’s F10 supplemented with 20% FBS and bFGF (0.025 μg/ml). For quiescence time point, cells were directly fixed with PFA 4% for 10min; for activation time points, cells were cultured for 24h and fixed with PFA 4%. Cells were incubated with rabbit anti-Dhx36 (Proteintech 13159-1-AP) primary antibody after blocking for 1 h at room temperature with 3% BSA (Sigma) in PBS. Cells were then washed with PBS and incubated with secondary antibody conjugated to AF488 or AF555 fluorochromes, and nuclei were stained with DAPI (Invitrogen) for 2 h at room temperature. After washing, Fluoromount (Thermo Fisher) was added to the wells of the chamber to preserve the fluorescence. For Pax7 staining on frozen muscle sections, an antigen retrieval step was performed before blocking through boiling samples in 0.01 M citric acid (pH 6.0) for 5 minutes in a microwave. After 4% BBBSA (4% IgG-free BSA in PBS; Jackson, ref: 001-000-162) blocking, the sections were further blocked with the Donkey anti-Mouse IgG (H+L) (1/100 in PBS; Jackson, ref: 115-007-003) for 30 minutes. The biotin conjugated anti-mouse IgG (1:500 in 4%BBBSA, Jackson, ref: 115-065-205) and Cy3-Streptavidin (1:1250 in 4%BBBSA, Jackson, ref: 016-160-084) were used as secondary antibodies. H&E staining on frozen muscle sections was performed as previously described[30]. All fluorescent images were captured with a fluorescence microscope (Leica). Primary antibodies used include: MyoD (1:100) (Santa Cruz Biotechnology) and Myogenin (1:200) (Santa Cruz Biotechnology); Pax7 (1:100) (Developmental Studies Hybridoma Bank) and MF20 (1:50) (Developmental Studies Hybridoma Bank); eMyHC (1:200) (Sigma-Aldrich) and laminin (1:800) (Sigma-Aldrich); Ki67 (1:200) (Thermo); MyoD (1:200) (Dako) for staining of muscle cryosections. Confocal images of isolated satellite cells were taken using a Zeiss LSM-780 confocal system with a Plan-Apochromat 63 × /1.4 NA oil. Acquisition was performed using Zeiss LSM software Zen Black. Images were slightly modified with ImageJ in which background was reduced using background subtraction and brightness and contrast were adjusted.

### CLIP-seq and data analysis

C2C12 cells in growth medium were crosslinked under UV and then lysed for immunoprecipitation using antibody against Dhx36 as previously described[60]. Retrieved RNPs were used for running PAGE gel and transferred to nitrocellulose membrane. RNA-protein complex with 15-20 kD larger than the protein MW were retrieved for library preparation and then sequenced with Illumina sequencing system. The first 6 nt of CLIP-seq raw reads were removed due to their low quality reported by FastQC[61], adaptors were further trimmed using Trimmomatic[62] with the minimal length threshold equal to 18 nt. After duplicates were removed, alignment was performed against mouse genome MM9 using Bowtie2[63] by allowing 2 insertions or deletions, 2 mismatches and only reads with greater than 90% identity were saved. Peak calling on the eligible reads was conducted via Piranha[64] using bin size parameter equal to 100 nt and significant threshold of 0.05. Reproducibility between biological replicates was examined by both Pearson Correlation Coefficient calculated using bam file coverages within 10kbp bin size from deepTools multiBamSummary module[65] and the proportion of peak overlapping using bedtools[66]. Genomic distribution of the peaks was annotated using mouse MM9 genome by HOMER[67] and in-house scripts.

To perform the de novo motif discovery on Dhx36 binding sites, MEME[31] was employed using -rna option, -nmotifs equal to 20 and 18 nt motif width, consistent with the length threshold used in CLIP-seq data preprocessing. The motif discovery on intronic binding sites was conducted on those sites not overlapping with intronic repeat elements. The most significant motifs and the corresponding E-values were reported. As for the identification of sequences similar with the 18 nt de novo motifs, FIMO[68] with “--norc --parse-genomic-coord” parameters was used.

To predict the potential rG4 forming sites in Dhx36 binding regions, we searched for the four subtypes defined as follows: G_3_L_1–7_, canonical rG4s with loop length between 1–7 nt (‘(G_3+_N_1–7_){3,}cG_3+_’, with *N* = A, U, C, or G); long loops, rG4s with any loop of length > 7 nt, up to 12 nt for lateral loops and 21 nt for the central loop (e.g., ‘G_3+_N_8–12_G_3+_N_1–7_G_3+_N_1–7_G_3+_’ or ‘G_3+_N_1–7_G_3+_N_13–_ 21G_3+_N_1–7_G_3+_’); bulges, rG4s with a bulge of 1–7 nt in one G tract, or multiple 1 nt bulges (e.g., ‘G_3+_N_1–9_G_3+_N_1–9_(GGH_1–7_G|GH_1–7_GG)N_1–9_G_3+_’ or ‘(GGHG|GHGG)N_1–9_ (GGHG|GHGG)N_1–9_G_3+_N_1–_9G_3+_’, with H = A, U, or C); 2 quartet, rG4s with four tracts of two consecutive Gs (’(G_2+_N_1–9_){3,}G_2+_’); G ≥ 50%, sequences that contain more than 50% G content and do not fall into the four previous categories; others, not in any previous category. When matching multiple categories, a region was assigned to the type with highest predicted stability, i.e. (from first to last), canonical rG4s, long loops, bulges and 2 quartet. When predicting rG4 formation in the 18 nt CLIP-seq motif enriched regions in Fig. 4K-M, only the rG4 sites overlapped with more than 9nt of the motif were used for the hierarchical assignment of rG4 subtypes described above.

To predict de novo structural motif, RNAshapes[34] tool was used to annotate the six generic shapes including Stems (S), Multiloops (M), Hairpins (H), Internal loops (I), dangling end (T) and dangling start (F) in RNA sequences. The structural motif logos were generated using WebLogo3 webserver[69].

To infer potential RBP binding in CDS sites, 8 nt motifs were compared with known RBP binding motifs from ATtRACT[70] database. Briefly, Position Weight Matrix (PWM) of all RBPs were downloaded from ATtRACT and converted to minimal MEME motif format by in-house scripts. Tomtom tool from the MEME suite was then utilized to compute the similarity between the 8 nt motifs and motifs in ATtRACT, with minimum overlapping threshold set to 5 nt and significance threshold equal to 3.0e-4.

To identify potential C/D box methylation guide snoRNAs, snoSCAN[35] was used with default parameters and settings. The Dhx36 intronic binding sites overlapped with neither annotated snoRNAs (information from HOMER software) nor intronic repeat elements (information downloaded from UCSC table browser[71]) were used as inputs for snoSCAN.

To identify the AU-rich element (ARE) in Dhx36 binding sites, we firstly searched for sequences matching three classes of conventional ARE patterns. The class one, defined as “several dispersed AUUUA motifs in an U-rich context”, included sequences matching the pattern of cluster 1 defined in [72]. We also added sequences matching ‘WWUUUWW’, ‘WWWUUUWWW’ and ‘WWWWUUUWWWW’ to class one, where W is A or U and these patterns were used by the ARE database AREsite2[73]. The definition of class two was identical with cluster 2,3,4,5 from [72], which includes AREs with two to five overlapping AUUUA motifs. The class three was defined as “U-rich regions” [33] so sequences with more than 5 continuous “U”s were collected. When matching multiple categories, the shorter AREs were removed. Finally, closely located sequences (within 5 nt) in classes one and three were merged together using bedtools merge, to produce the final ARE set.

To calculate the Minimum Free Energies (MFEs) of RNA secondary structures, RNAfold program from ViennaRNA 2.4.10[74] was used. The MFEs of dsRNA secondary structures were calculated at 37 °C with parameters “--MEA -p0 -d2 --noLP” and named as 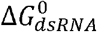 while MFEs for rG4 structures were calculated using the following formula:

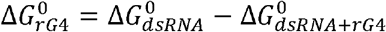

where 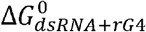 were the MFEs calculated when considering rG4 formation into RNAfold prediction. Both MFE values were normalized by the input sequence length and the types of input sequence were the 5’UTR regions and the ARE sites with a 30 nt extension on both sides. For all the MFE comparisons, we randomly selected 1000 genes without Dhx36 bindings and showed no TE or mRNA changes as the background set.

### Polysome profiling and data analysis

Ctrl or *Dhx36* KO C2C12 cells were cultured in growth medium. Prior to lysis, cells were treated with cycloheximide (100 μg/ml) for 10 min at 37°C. Cells were then washed with ice□cold PBS supplemented with 100 μg/ml cycloheximide and further lysed in 300 μl of lysis buffer (10 mM HEPES pH 7.4, 150 mM KCl, 10 mM MgCl2, 1% NP□40, 0.5 mM DTT, 100 μg/ml cycloheximide). After lysing the cells by passing eight times through 26□gauge needle, the nuclei and the membrane debris were removed by centrifugation (15,682 g, 10 min, 4°C). The supernatant was then layered onto a 10□ml linear sucrose gradient (10–50% [w/v], supplemented with 10 mM HEPES pH 7.4, 150 mM KCl, 10 mM MgCl2, 0.5 mM DTT, 100 μg/ml cycloheximide) and centrifuged (160,000 g, 120 min, 4°C) in an SW41Ti rotor (Beckman). After centrifugation the sediment in the tube went through a UV detector at the wavelength of 254nm and simultaneously the required fractions (>=5 ribosomes, polysome) were collected and polysome-associated RNAs were extracted by Trizol-LS. TruSeq Stranded Total RNA libraries were prepared with 500 ng RNA according to the manufacturer’s protocol (Illumina). The libraries were sequenced in 2 × 100 nt manner on HiSeq 2000 platform (Illumina).

The sequenced total cytoplasmic mRNA and polysome-associated RNA libraries were quantified following the RNA-seq processing procedures. Briefly, after the adaptor trimming, quality filtering and duplication removal using in-house scripts[75], the sequenced fragments were mapped to reference mouse genome (MM9) using TopHat2[76]. Cufflinks[77] was then used to derive the Fragments Per Kilobase per Million (FPKM) values. TE values were then calculated as the ratio between the abundance of polysome-associated RNAs (FPKM-polysome) and total cytoplasmic mRNAs (FPKM-total) in Ctrl and *Dhx36* KO conditions. The TE values in KO were further divided by TEs in Ctrl, generating log2(ΔTE)). Only those genes with TE up- or down-regulated in both replicates, were used for further analysis with their averaged FPKM-polysome and FPKM-total values. When intercepting with CLIP-seq data to dissect the effect of Dhx36 binding on TE and total mRNA abundance, transcripts with averaged FPKM-total values larger than 1 in either Ctrl or KO condition were used.

### Gene Ontology analysis

ClusterProfiler [78] was used for the Gene Ontology (GO) analysis with Entrez gene IDs converted from DAVID[79] tool as inputs. The adjusted P or P. values were reported with the GO terms.

### Statistical analysis

The statistical significance of experimental data was calculated by the Student’s t-test (two-sided). *P<0.05, **P<0.01, ***P<0.001 and n.s.: not significant (p>=0.05). The statistical significance for the assays done with SCs from the same mouse with different treatment was calculated by the Student’s t-test (paired). *P<0.05, **P<0.01, ***P<0.001 and n.s.: not significant (p>=0.05). The statistical significance of MFEs between groups was assessed using one-tailed Mann-Whitney test. ***P<0.001, and n.s.: not significant (p>=0.05).

## Data availability

CLIP-seq and Polysome profiling data used in this study have been deposited in Gene Expression Omnibus (GEO) database under the accession codes GSE151124.

## Code availability

The code for rG4 site and AU-rich element identification has been deposited in a Github repository (https://github.com/jieyuanCUHK/DHX36_paper).

## Author Contributions

X.C. and H.W. designed the experiments; X.C. and G.X. conducted the experiments; Y.Z., L.H., Y.L. provided technical supports; J.Y. analyzed the NGS data; S.C.S., J.I., C.M. and E.P. performed Dhx36 cKO mice and part of iKO mice analyses; W.W. and W.C. contributed to Polysome profiling; X.M conducted RTS gel assay; M.I.U. conducted ThT and NMM staining; D.W., L.W. and Y.X. contributed to CLIP-seq; Y.N. generated the *Dhx36^fl/fl^* mice; C.K.K supervised rG4 analyses. H.S. supervised computational analyses. P.M.C. supervised Dhx36 cKO mice and part of iKO mice analyses. X.C., J.Y. and H.W. wrote the paper.

## Acknowledgements

We thank Prof. Zhongzhou Yang for his generous sharing of the *Dhx36^fl/fl^* mice and the pEGFP-N1 plasmid. This work was supported by General Research Funds (GRF) from the Research Grants Council (RGC) of the Hong Kong Special Administrative Region (14115319, 14100018, 14133016 and 14106117 to H.W.; 14116918 and 14120619 to H.S.); the National Natural Science Foundation of China (NSFC) to H.W. (Project code: 31871304), NSFC/RGC Joint Research Scheme to H.S. (Project code: N_CUHK 413/18); Focused Innovations Scheme: Scheme B to H.S. [Project Code: 1907307]; Natural Science Foundation of Guangdong Province to X.C. [Project code: 2019A1515010670]. Work in PMC laboratory was supported by Spanish Ministry of Science, Innovation and Universities, Spain (grants RTI2018-096068-B-I00 and SAF 2015-70270-REDT, a María de Maeztu Unit of Excellence award to UPF [MDM-2014-0370], and a Severo Ochoa Center of Excellence award to the CNIC [SEV-2015-0505]), ERC-2016-AdG-741966, La Caixa-HEALTH (HR17-00040), MDA, UPGRADE-H2020-825825, AFM and DPP-E. SC is recipient of a FI fellowship from AGAUR.

